# RNA Polymerase Backtracking Promotes Type III-A CRISPR DNA Targeting

**DOI:** 10.64898/2026.11.04.764256

**Authors:** Angel J. Hernandez Viera, Naomi Gross, Patricia Q. Tran, Omar Shahin, James C. Kosmopoulos, Karthik Anantharaman, Charlie Y. Mo

**Affiliations:** Department of Bacteriology, University of Wisconsin-Madison; Microbiology Doctoral Training Program, University of Wisconsin-Madison

## Abstract

Type III CRISPR-Cas systems provide transcription-dependent immunity against mobile genetic elements, yet the mechanism underlying their DNA targeting remains unclear. Although the type III effector Cas10-Csm functions as a single-stranded DNase *in vitro*, type III systems eliminate double-stranded DNA targets *in vivo*, suggesting that transcription generates critical DNA substrates. Here, we show that conserved RNA polymerase anti-backtracking factors, such as GreA, selectively inhibit DNA-targeting of type III immunity. GreA overexpression abolishes protection against both phages and conjugative plasmids, whereas mutations that disrupt GreA anti-backtracking activity eliminate this effect. In contrast, immunity mediated by the type II CRISPR-Cas system or the auxiliary type III RNase Csm6 remain unaffected, demonstrating that GreA specifically suppresses Cas10-dependent DNA interference. These findings identify RNA polymerase backtracking as a key determinant of Cas10-Csm activity and support a model in which arrested transcription complexes generate substrates for DNA degradation, opening up new anti-CRISPR strategies against type III.

## Introduction

CRISPR-Cas systems are a form of prokaryotic immunity that protects bacteria and archaea from foreign nucleic acids, such as plasmids, and bacteriophages. CRISPR-Cas systems create a memory of past infections by acquiring short sequences of invading DNA, called spacers^1,2^. These spacers are transcribed and processed into CRISPR RNAs (crRNA) which then form an effector complex with CRISPR-Associated (Cas) proteins^2^. After finding a target sequence with complementary base pair binding, the CRISPR effector complex activates nucleases, which cut and degrade the invading nucleic acid, thus providing immunity against the parasitic element.

Of the seven different types of CRISPR-Cas systems (types I through VII) identified to date, the type III system is the most functionally elaborate and evolutionarily ancient^3, 4^. The Cas effector of type III-A CRISPR is a multiprotein complex, called Cas10-Csm, consisting of the signature protein Cas10, four additional subunits (Csm2-5), and the CRISPR-guide RNA (crRNA). Type III-A CRISPR defense is dependent on transcription of the target sequence, since Cas10-Csm can only bind to complementary RNA^5,6^. Cas10-Csm binding to the target transcript triggers the activities of two domains within the Cas10 protein: a single-stranded DNase activity, conferred by the HD domain, that degrades single-stranded DNA of any sequence; and a cyclase activity, conferred by the Palm domain, that synthetizes cyclic oligoadenylates (cOAs) from ATP ^6–8^. cOAs serve as a secondary messenger that activates ancillary proteins with a CRISPR-associated Rossman fold (CARF) domains, such as the nonspecific RNase, Csm6 ^9,10^. Finally, target recognition activates a sequence-specific RNase in the Csm3 subunits of the Cas10-Csm complex, which degrades the bound RNA target, resulting in inactivation of all enzymatic activities. Csm3 activity is not required for phage or plasmid interference^11,12^, but likely limits the damage to the host that would otherwise be caused by the uncontrolled non-specific ssDNase and RNase activities of Cas10 and Csm6, respectively.

A central unresolved question in type III CRISPR biology is how the Cas10-Csm effector complex mediates DNA degradation *in vivo*. Biochemical studies have shown that purified Cas10-Csm cleaves only single-stranded DNA (ssDNA) substrates^6,13,14^, yet type III immunity in cells results in the rapid destruction of double-stranded DNA targets, including plasmids and phage genomes^10,12^. How type III systems bridge this apparent discrepancy remains unknown. Accessible ssDNA is relatively rare in bacterial cells but occurs transiently at replication forks and transcription bubbles. Because type III immunity requires transcription of the target sequence, a prevailing model proposes that Cas10-Csm exploits the exposed ssDNA within the transcription bubble generated by RNA polymerase (RNAP). However, the evidence supporting this model remains incomplete and, in some cases, contradictory. *In vitro* co-transcriptional cleavage assays suggest that Cas10 targets the exposed non-template DNA strand within the transcription elongation complex (TEC)^13,15^, consistent with transcription-dependent DNA degradation. In contrast, Liu *et al* showed that, although tethering of the type III effector complex to the transcription bubble can occur, there is no direct interaction to the RNAP or ssDNA in the transcription bubble^16^. Furthermore, they did not observe DNA degradation at the transcription bubble^16^, raising questions about how the effector complex accesses target DNA within actively transcribing complexes. These observations suggest that transcription is essential for type III-mediated DNA degradation, yet the mechanistic relationship between transcription dynamics and Cas10 activity remains poorly understood. Here, we investigate how transcription influences DNA targeting and degradation by the Cas10-Csm complex, with the goal of defining the molecular basis of type III CRISPR immunity inside cells.

RNA polymerase (RNAP) does not transcribe DNA in a continuous manner but instead undergoes pausing and backtracking during elongation. During backtracking, RNAP reverse-translocates and displaces the 3’ end of the nascent transcript from the active site into the secondary channel blocking further transcription elongation^17^. Backtracking can be triggered by a variety of obstacles, including DNA damage^18^, nucleotide misincorporation into the nascent transcript^19^, and collisions with DNA-bound proteins or the replisome^20^. Persistent backtracked RNAP complexes threaten genome integrity because they obstruct DNA replication and other essential genome maintenance processes^17^. To mitigate these conflicts, both prokaryotic and eukaryotic cells encode anti-backtracking factors that reactivate stalled transcription complexes and restore productive elongation^21^.

Here, we show that overexpression of anti-backtracking factors, such as GreA, selectively inhibits Cas10-Csm-mediated DNA targeting of type III immunity, resulting in a loss of protection against both phages and conjugative plasmids. This inhibitory effect depends on the anti-backtracking activity of GreA, as it is eliminated by mutations that disrupt GreA function. Moreover, GreA-mediated inhibition is bypassed under conditions that promote activation of the Csm6 RNase, indicating that interference is specific to the DNA-targeting branch of type III immunity. Our results identify RNAP backtracking as a critical determinant of Cas10-Csm activity and suggest that transcriptional roadblocks or arrested elongation complexes may provide key substrates for DNA degradation by type III systems. More broadly, our findings demonstrate that manipulating transcription dynamics can suppress type III CRISPR immunity and reveal a previously unrecognized mechanism for antagonizing CRISPR defense. To our knowledge, this study provides the first evidence that perturbation of RNAP backtracking can function as an anti-CRISPR strategy against type III systems.

### RNAP anti-backtracking inhibits type III-A CRISPR anti-phage immunity in *Staphylococcus aureus*

Because type III CRISPR immunity is mechanistically linked to transcription, we investigated whether reducing RNAP backtracking affects defense against bacteriophage infection. In many bacterial species, backtracked RNAP complexes are rescued by the transcription factor GreA, which stimulates cleavage of the extruded RNA transcript and realigns its 3′ end within the RNAP active site, thereby restoring productive elongation^22,23^. Overexpression of GreA has been shown to reduce RNAP backtracking^24^. To test whether diminished RNAP backtracking alters type III immunity, we cloned the *S. aureus greA* gene into an inducible expression vector (pGreA) and introduced either pGreA or an empty-vector control (pEV) into *S. aureus* RN4220. We then established type III immunity by introducing a plasmid-borne *S. epidermidis* type III-A CRISPR-Cas system (pCRISPR)^25^ carrying a spacer targeting the early phage gene *gp14* from the lytic staphylococcal phage ΦNM1γ6.

We challenged these strains with ΦNM1γ6 in plaque formation assays. As expected, RN4220 carrying pEV exhibited robust type III immunity, preventing the formation of clear viral plaques (**Figure 1A**). In contrast, GreA overexpression increased phage plaquing efficiency by approximately three orders of magnitude. Consistent with this result, GreA overexpression increased the average burst size of ΦNM1γ6 by approximately ten-fold relative to the EV control (**Figure 1B**) and significantly impaired bacterial growth during phage infection (**Figure 1C**). To determine whether this effect was specific to type III immunity, we repeated burst-size, and growth assays using a type II CRISPR-Cas system (Cas9) targeting the same phage. In this context, GreA overexpression had no detectable effect on immunity (**Figure S1A and S1B**), indicating that GreA specifically interferes with Cas10-Csm-mediated defense rather than CRISPR immunity in general.

**Figure 1.**
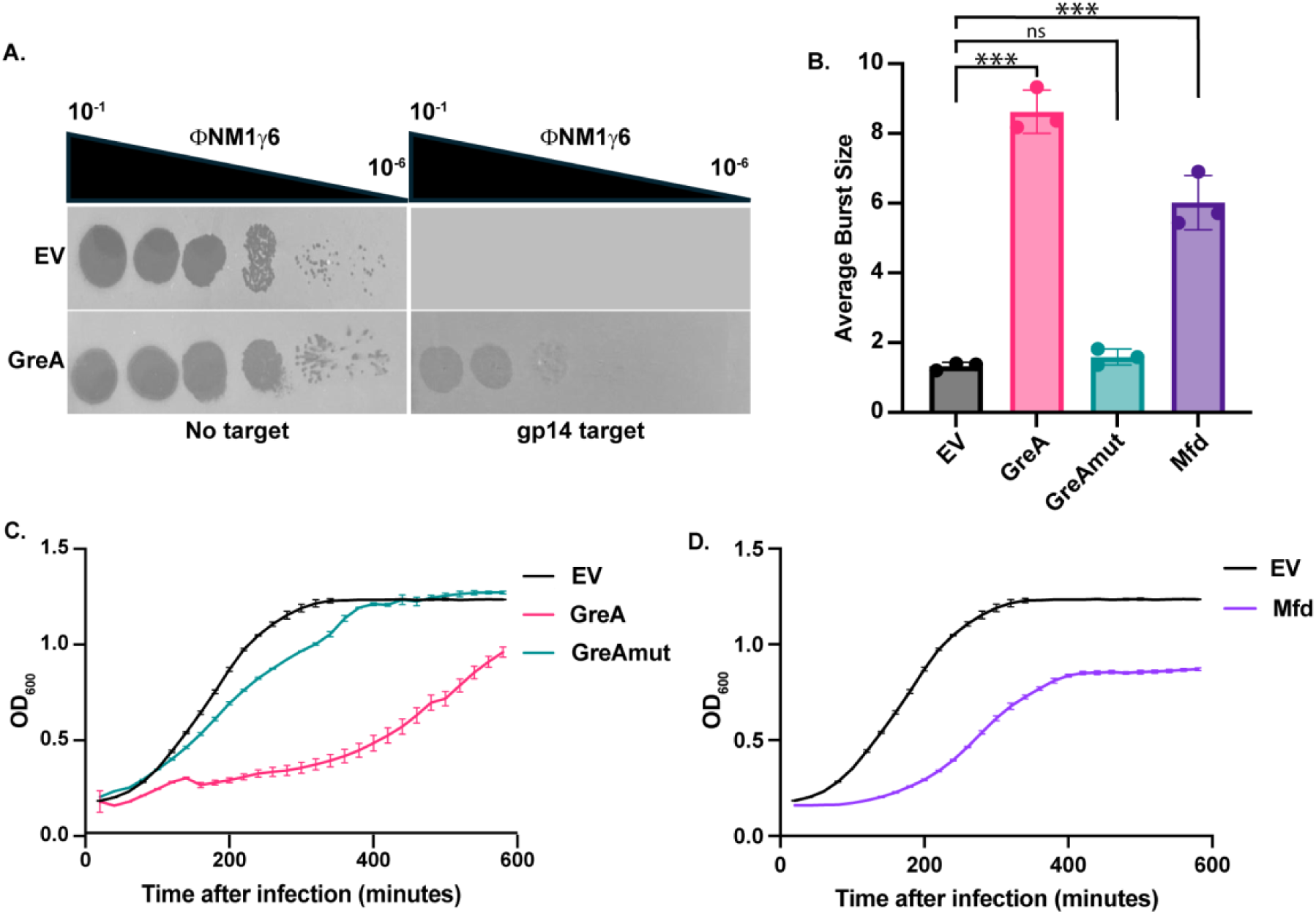
RNA polymerase anti-backtracking factors inhibit type III-A CRISPR anti-phage defense against ΦNM1γ6. **A.** Serial ten-fold dilutions of ΦNM1γ6 plated on *S. aureus* RN4220 containing type III-A CRISPR system with or without a spacer targeting the *gp14* gene of ΦNM1γ6. Bacterial cells also contain the GreA overexpression plasmid or the empty vector (EV). **B.** Average burst size of ΦNM1γ6 in RN4220 co-transformed with a type III-A CRISPR system targeting *gp14* and an empty (EV), GreA, GreAmut, or Mfd overexpression plasmid. Mean ± SD of 3 replicates is reported. **C.** Growth curves of RN4220 with the type III-A CRISPR targeting *gp14* in the presence of ΦNM1γ6 at an MOI of 0.1. Cells are also co-transformed with the GreA or GreAmut overexpression plasmid, or the empty vector (EV). Mean ± SD of 3 replicates is reported. **D.** Same as **C** but with cells carrying an Mfd overexpression plasmid or the empty vector (EV). Where applicable, statistics were performed with an unpaired, two-sided t-test. ns: p>0.05; *: p<0.05; **: p<0.01; ***: p<0.001

We next asked whether the observed phenotype depended on the anti-backtracking activity of GreA. To test this possibility, we generated a catalytic mutant allele, *greAD44S,E47S* (GreAmut), containing substitutions in two conserved acidic residues required for rescue of backtracked RNAP complexes^26^. In contrast to wild-type GreA, expression of GreAmut had no effect on phage plaquing efficiency (**Figure 1A**), burst size (**Figure 1B**), or bacterial growth (**Figure 1C**) relative to the EV control, demonstrating that the anti-backtracking activity of GreA is required to inhibit type III immunity. To independently evaluate the role of RNAP backtracking, we overexpressed the transcription-repair coupling factor Mfd, which resolves stalled transcription complexes through a mechanism distinct from that of GreA^27^. Similar to GreA, Mfd overexpression significantly reduced type III-mediated protection against phage infection (**Figure 1B and 1D**). Because RNAP backtracking is frequently associated with R-loop formation^28^, we also examined the effect of overexpressing RNaseHIII, which selectively degrades R-loops^29^. In contrast to GreA and Mfd, RNaseHIII overexpression had no detectable effect on type III immunity (**Figure S2A and S2B**).

Because reducing RNAP backtracking could change global transcriptional programs, we sought to determine whether the inhibitory effect of GreA on type III immunity was an indirect consequence of changes in host transcript levels. We performed RNA-seq on RN4220 carrying either pGreA or the empty-vector control and conducted differential expression analysis across the transcriptome. As expected, GreA overexpression did not significantly alter transcript levels of the plasmid-borne type III CRISPR-Cas locus, indicating that the loss of immunity cannot be explained by reduced CRISPR-Cas expression (**Supplemental File 1**). Outside of *greA,* the only overexpressed gene was *udk,* which is encoded immediately downstream of *greA.* We also observed a group of genes being downregulated when GreA is overexpressed: *cwrA*^30^, which responds to cell wall stress and when deleted causes loss of cell wall integrity; *sgtB* (also known as *mgt)*^31^, a non-essential glycosyltransferase; *vraX,* a secreted protein that is upregulated under cell wall stress^32,33^; as well as a group of small proteins annotated as hypothetical proteins (**Figure S3A**). As lytic phages need to bind, penetrate, and rupture the bacterial cell wall during their life cycle, it is formally possible that these genes could affect the cell wall integrity and therefore the phage infectious cycle. However, we think this scenario is unlikely: in the absence of CRISPR targeting, GreA overexpression only resulted in a modest increase in ΦNM1γ6 burst size of less than two-fold (**Figure S3B**) and did not measurably increase phage plaquing efficiency (**Figure 1A**). Furthermore, type II immunity was not affected by GreA overexpression.

Although the increase in phage production was less that two-fold, it was nonetheless statistically significant (**Figure S3B**). We therefore asked whether this small increase in phage production could account for the growth defect observed during infection when GreA was overexpressed in the presence of type III immunity. To address this possibility, we compared bacterial growth following infection at multiplicities of infection (MOIs) of 0.1 and 1, reasoning that the higher MOI would approximate the increased phage burden observed in GreA-overexpressing cells. Despite the ten-fold difference in viral input, no difference in bacterial growth was observed, indicating that enhanced phage production alone is insufficient to explain the GreA-dependent loss of protection (**Figure S3C)**. We also developed a self-targeting system in which the targeted genome is the bacterial chromosome; in this system, our measurements do not depend on the targeted genomes going through the cell wall (see ‘Anti-backtracking of RNAP reduces DNA degradation caused by type III immunity’ section below).

Lastly, we accounted for the possibility that GreA overexpression might generate more phage escapers against type III targeting. However, consistent with previous studies^34^, we were unable to detect phage escapers during type III immunity in either the presence or absence of GreA overexpression. In contrast, escape mutants were readily recovered from strains carrying a type II CRISPR-Cas system targeting the same phage, as expected^34^. Importantly, GreA overexpression did not alter the frequency of type II escape mutants (**Figure S4A**), further arguing against a mechanism that enhances phage escaper formation. In addition, we found that the inhibition of immunity caused by GreA overexpression is independent of the SOS response (**Figure S4B)**, which induces mutagenic DNA repair and is activated by type III CRISPR immunity^8^. Together, these results eliminate several indirect explanations for the loss of type III immunity, including altered CRISPR-Cas expression, enhanced phage replication and phage escape. Instead, they support a model in which RNAP backtracking directly promotes Cas10-Csm-mediated defense and wherein reducing backtracking suppresses type III immunity by interfering with the DNA-targeting activity of the effector complex.

### RNAP anti-backtracking inhibits ssDNase-dependent type III immunity during phage infection

To determine how GreA inhibits type III immunity, we directly monitored phage DNA levels following infection. We designed qPCR primers targeting two loci within the ΦNM1γ6 genome: the CRISPR-targeted gene, *gp14*, and *gp43*, located approximately 21 kb away from the target site. We then quantified the abundance of both loci in infected cells over time. Consistent with previous reports^10^, phage DNA levels declined in cells carrying the type III CRISPR-Cas system and empty vector (EV), with both *gp14* and *gp43* decreasing within minutes of infection (**Figure 2A: left panel)**. This reduction is indicative of Cas10-Csm-mediated degradation of the phage genome. In contrast, with pGreA, the abundance of both loci increased throughout the course of infection, reaching approximately 50-fold higher levels by 90 minutes post-infection. Notably, the increase was observed for both the targeted locus (*gp14*) and a distal region (*gp43*) of the genome, suggesting that phage DNA replication proceeded despite CRISPR targeting.

**Figure 2.**
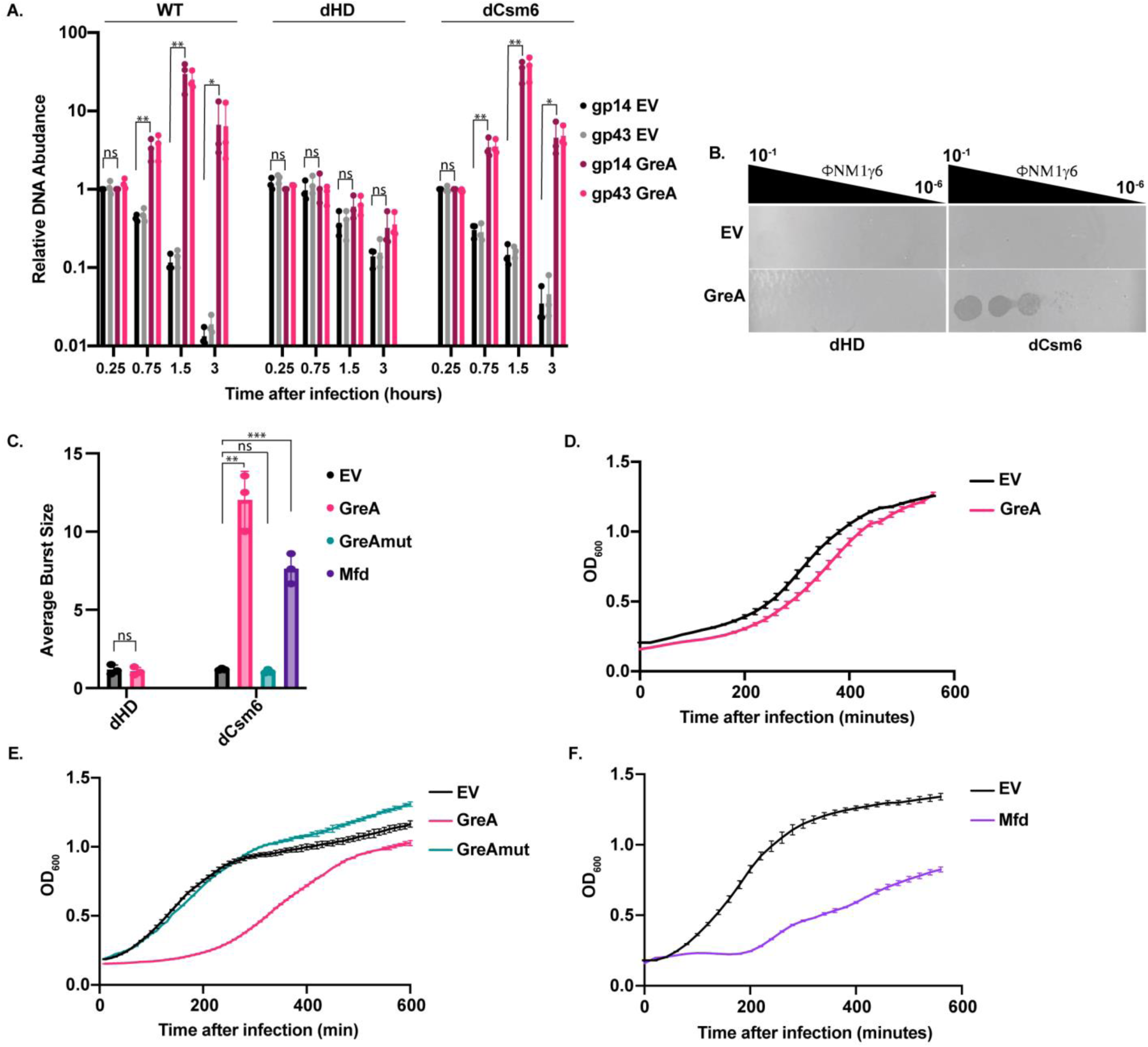
RNA polymerase anti-backtracking factors inhibit type III-A CRISPR ssDNase-dependent anti-phage defense. **A.** qPCR of the *gp14* or *gp43* gene of ΦNM1γ6 using total DNA extracted after phage infection of RN4220 carrying wild-type (WT), HD domain-deactivated (dHD), or Csm6 deactivated (dCsm6) type III-A CRISPR. All systems target the *gp14* gene. The cells also carry a GreA overexpression plasmid or the empty vector (EV). Mean ± SD of 3 replicates is reported. **B.** Serial ten-fold dilutions of ΦNM1γ6 plated on *S. aureus* RN4220 with dHD or dCsm6 type III-A CRISPR system targeting the phage *gp14* gene. Cells also contain a GreA overexpression plasmid or the empty vector (EV). **C.** Average burst size of ΦNM1γ6 in RN4220 containing a HD domain-deactivated (dHD) or a Csm6-deactivated (dCsm6) type III-A CRISPR targeting the phage *gp14* gene, and a GreA, GreAmut, or Mfd overexpression plasmid, as well as the empty vector (EV). Mean ± SD of 3 replicates is reported. **D.** RN4220 growth curve containing a HD domain-deactivated type III-A CRISPR targeting the phage *gp14* gene in the presence of ΦNM1γ6 at an MOI of 0.1. Cells also contain a GreA overexpression plasmid or the empty vector (EV). Mean ± SD of 3 replicates is reported. **E.** Same as **D**, but with cells containing a Csm6-deactivated type III-A CRISPR and a GreA or GreAmut overexpression plasmid, or the empty vector (EV). **F.** Same as (D) but with cells carrying an Mfd overexpression plasmid or the empty vector (EV). Where applicable, statistics were performed with an unpaired, two-sided t-test. ns: p>0.05; *: p<0.05; **: p<0.01; ***: p<0.001

The type III-A CRISPR-Cas system protects against phage infection through two mechanistically distinct pathways: sequence-independent ssDNA degradation mediated by the HD nuclease domain of Cas10 and sequence-independent RNA degradation mediated by the accessory nuclease Csm6.^6,10^ We therefore asked whether GreA overexpression preferentially interferes with Cas10-dependent DNA targeting or Csm6-mediated RNA targeting. Importantly, inactivation of either nuclease does not abolish immunity against phages but instead shifts the mechanism by which defense is achieved^11,10^.

To dissect the branch of immunity affected by GreA, we used type III variants that rely exclusively on either Csm6 RNase activity (dHD) or Cas10 HD nuclease activity (dCsm6). These systems were introduced into strains carrying either pGreA or pEV, and phage DNA levels were quantified following infection using the qPCR assay described above. In the pEV background, the dCsm6 mutant exhibited a decline in phage DNA abundance following infection, consistent with active Cas10-mediated DNA degradation (**Figure 2A: right panel)**. In contrast, phage DNA levels declined more slowly in the dHD mutant (**Figure 2A: center panel)**. This phenotype is consistent with the established mechanism of Csm6-dependent immunity: in the absence of the Cas10 HD nuclease, phage genomes are not extensively degraded but are nevertheless unable to replicate efficiently because persistent Csm6 activation causes widespread RNA destruction. Thus, although both systems remain protective, they differ markedly in their effects on phage DNA dynamics.

Strikingly, GreA overexpression selectively affected the HD-dependent pathway. In the dCsm6 background, GreA overexpression caused phage DNA levels to increase over time, reaching an approximately 50-fold higher abundance than the EV control, a phenotype comparable to that observed with the wild-type type III system (**Figure 2A: right panel)**. In contrast, GreA overexpression had no significant effect on phage DNA abundance in the dHD background (**Figure 2A: center panel)**. These findings indicate that GreA specifically disrupts Cas10-mediated DNA targeting while leaving Csm6-dependent immunity largely intact.

This selective inhibition was also evident across multiple independent measures of host protection. GreA overexpression significantly increased both plaquing efficiency and phage burst size in strains carrying the dCsm6 system but had little or no effect on these parameters in strains carrying the dHD system (**Figure 2B and 2C)**. Likewise, GreA overexpression impaired bacterial growth during phage infection in the dCsm6 background but did not measurably affect growth of strains protected solely by Csm6-mediated immunity (**Figure 2D and 2E)**. Importantly, these effects required the anti-backtracking activity of GreA, as the catalytically inactive GreA mutant (GreAmut) failed to increase burst size or compromise host growth in the dCsm6 strain (**Figure 2C and 2E)**. We also obtained similar results with Mfd: overexpression of Mfd increased phage burst size and reduced bacterial growth during infection in the dCsm6 background, mirroring the effects of GreA (**Figure 2C and 2F)**. The concordance between these two mechanistically distinct anti-backtracking factors strongly supports the conclusion that RNAP backtracking promotes Cas10-dependent immunity. We do note that since GreA and Mfd did not affect immunity conferred by Csm6, it remains unclear how GreA permits phage propagation in the strain containing wild-type type III CRISPR, which has both HD and Csm6 activities (**Figure 2A, 2B, and 2C)**. We speculate that that reduced backtracking might have a small inhibitory effect on Csm6-based immunity that we are not capturing in our assays.

Together, these findings demonstrate that reducing RNAP backtracking impairs the DNA-targeting branch of type III CRISPR immunity. Because GreA and Mfd inhibit protection only when defense relies on the Cas10 HD nuclease, but not when immunity is mediated by Csm6, we conclude that RNAP backtracking is specifically required for efficient Cas10-dependent DNA degradation. These results further suggest that backtracked transcription complexes may serve as critical substrates or intermediates that enable Cas10-Csm to access and destroy target DNA during infection.

### Effect of GreA anti-backtracking activity on type III immunity targeting late genes of the phage

A unique feature of type III CRISPR immunity is that the mechanism of defense depends on which phage gene is targeted. Previous studies have shown that targeting early phage transcripts primarily relies on the HD nuclease activity of Cas10^11^, whereas targeting late phage transcripts requires the Csm6 for robust antiphage defense^11,10^; under these conditions, HD-mediated DNA degradation plays a more limited role^11^. Because our results indicated that GreA overexpression selectively inhibits Cas10-dependent immunity, we hypothesized that GreA would have little effect on type III defense when the targeted transcript is expressed late during the phage infection cycle.

To test this hypothesis, we used pCRISPR constructs targeting the late phage genes *gp43* and *gp59* and evaluated the impact of GreA overexpression on antiphage immunity. Consistent with our prediction, GreA overexpression did not measurably increase phage propagation when late genes were targeted, as assessed by either plaque formation or burst-size assays (**Figure 3A, 3B, and S5A**). However, we observed a modest reduction in bacterial growth relative to strains targeting the early gene *gp14* (**Figure S5B)**. Given that GreA inhibits Cas10-dependent DNA degradation, we propose that phage genomes are not efficiently cleared from the bacterial population under these conditions. Persistent phage DNA would allow continued transcription of the target gene, resulting in prolonged activation of Csm6 and increased collateral RNA degradation, which likely contributes to the observed reduction in bacterial growth.

**Figure 3.**
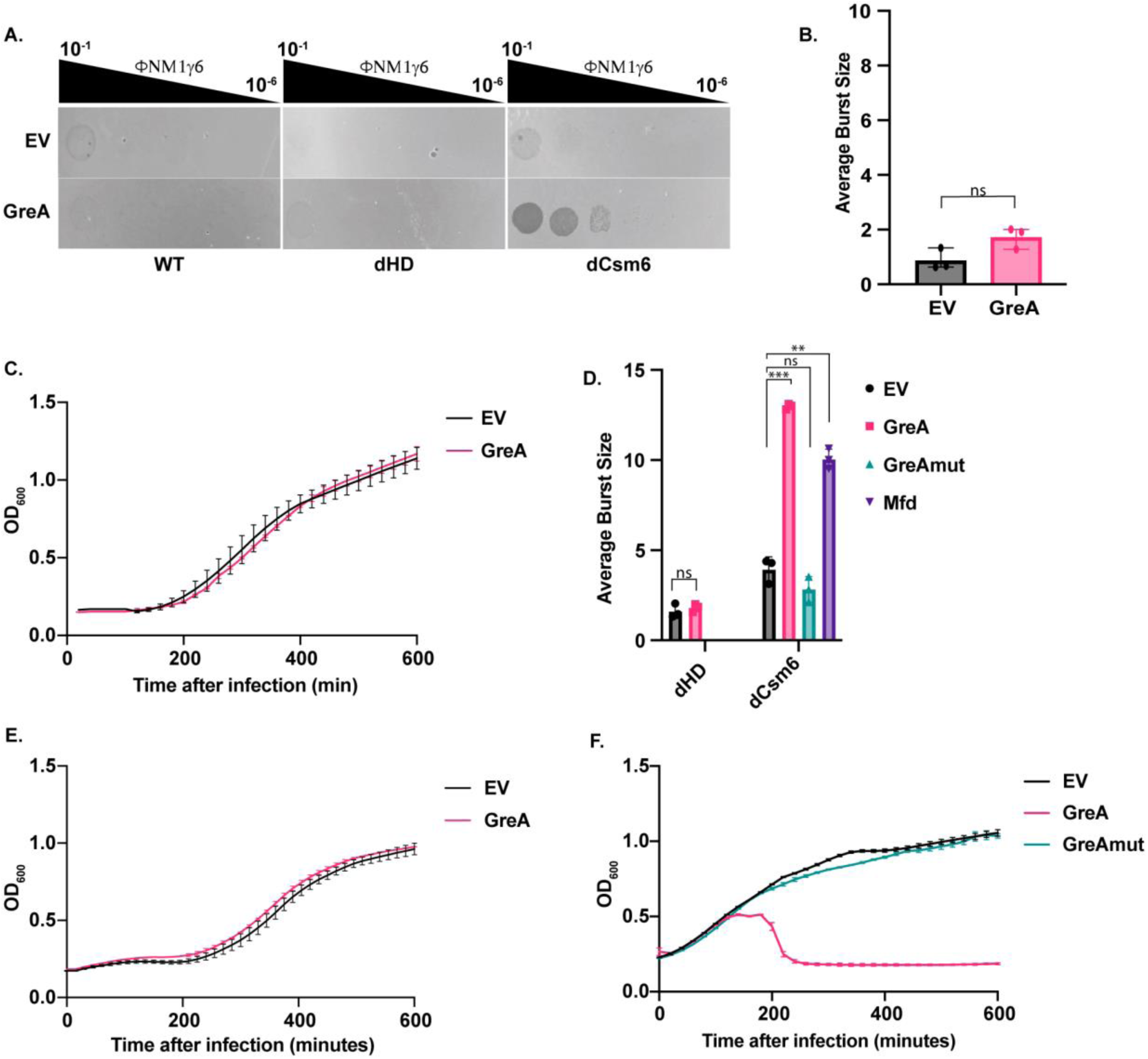
Anti-backtracking factors do not inhibit type III-A CRISPR anti-phage defense when targeting a late gene. **A.** Serial ten-fold dilutions of ΦNM1γ6 plated on *S. aureus* RN4220 carrying a wild-type (WT), HD domain-deactivated (dHD), or Csm6 deactivated (dCsm6) type III-A CRISPR system targeting the *gp43* gene. Cells also contain a GreA overexpression plasmid or the empty vector (EV). **B.** Average burst size of ΦNM1γ6 in RN4220 containing type III-A CRISPR targeting the phage *gp43* late gene and a GreA overexpression plasmid, or the empty vector (EV). Mean ± SD of 3 replicates is reported. **C.** RN4220 growth curve containing type III-A CRISPR targeting the phage *gp43* late gene in the presence of ΦNM1γ6 at an MOI of 0.1. Cells also contain a GreA overexpression plasmid, or the empty vector (EV). Mean ± SD of 3 replicates is reported. **D.** Average burst size of ΦNM1γ6 in RN4220 containing a HD domain-deactivated (dHD) or a Csm6-deactivated (dCsm6) type III-A CRISPR targeting *gp43*, and a GreA, GreAmut, or Mfd overexpression plasmid, as well as the empty vector (EV). Mean ± SD of 3 replicates is reported. **E.** Same as **C**, but with cells containing a HD domain-deactivated type III-A CRISPR. **F.** Growth curve of RN4220 with a Csm6-deactivated type III-A CRISPR targeting the phage *gp43* late gene in the presence of ΦNM1γ6 at an MOI of 0.01. Cells also contain a GreA, or GreAmut overexpression plasmid, or the empty vector (EV). Mean ± SD of 3 replicates is reported. Where applicable, statistics were performed with an unpaired, two-sided t-test. ns: p>0.05; *: p<0.05; **: p<0.01; ***: p<0.001

We next examined the effects of GreA overexpression on the dHD and dCsm6 mutants targeting the late gene *gp43*. Consistent with the results obtained for early-gene targeting, GreA had no detectable effect on phage propagation or bacterial growth in the dHD strain, where immunity depends entirely on Csm6. In contrast, because Csm6 is required for efficient defense against late-gene targets, the dCsm6 strain exhibited sensitivity to infection, we therefore performed these experiments at an MOI of 0.01, under which the strain remained viable and the effects of GreA could be evaluated. Under these conditions, GreA overexpression, but not GreAmut overexpression, increased phage burst size and further reduced bacterial growth relative to the pEV control. Thus, as observed with early-gene targets, reducing RNAP backtracking compromises Cas10-dependent immunity when targeting late phage genes.

Lastly, we asked whether these findings extended beyond ΦNM1γ6. To address this question, we examined the effect of GreA overexpression on type III-A defense against the staphylococcal phage Φ12. The same pattern was observed: GreA overexpression impaired immunity when an early phage gene was targeted but had little effect when a late gene was targeted (**Figure S5 C-E**). Together, these results indicate that the inhibitory effects of GreA are not phage-specific and further support the conclusion that RNAP backtracking is specifically required for Cas10-mediated DNA degradation. More broadly, these findings reinforce a model in which anti-backtracking factors selectively suppress the DNA-targeting branch of type III immunity while leaving Csm6-dependent defense largely unaffected.

### RNA polymerase backtracking promotes Cas10 ssDNase-mediated anti-plasmid immunity

The type III-A CRISPR-Cas system from *S. epidermidis* RP62a naturally carries a spacer targeting the *nickase* (*nes*) gene of the conjugative plasmid pG0400, which confers mupirocin resistance^25^. We therefore asked whether reducing RNAP backtracking through GreA overexpression could impair type III-mediated anti-plasmid defense. To test this possibility, we constructed *S. aureus* strains carrying either pGreA or the empty-vector control together with a pCRISPR plasmid containing the native spacer 1 targeting pG0400 (spc1). We then measured plasmid transfer by conjugating pG0400 into these recipient strains and quantifying conjugation efficiency as the ratio of mupirocin-resistant transconjugants to total colony-forming units.

As expected, type III CRISPR targeting substantially reduced conjugation efficiency relative to strains lacking targeting type III CRISPR (no spacer) (**Figure 4A)**. Unlike against phages, however, GreA overexpression did not significantly alter conjugation efficiency in the wild-type type III-positive backgrounds (**Figure 4A)**. This result was unexpected because previous work by Rostøl and Marraffini demonstrated that anti-plasmid immunity against pG0400 depends on the Cas10 HD nuclease domain^12^, the same pathway inhibited by GreA in our phage experiments. We also repeated this experiment but using a CRISPR system with Csm6 deactivated (dCsm6) to investigate the effect of GreA overexpression on the ssDNase-based immunity; however, as Rostøl and Marraffini showed in their study, we observed that Csm6 is required for plasmid immunity in this context. Notably, that study also showed that the native spacer targets a weakly expressed antisense transcript within the *nes* locus. We therefore posited that low target transcription might limit the impact of GreA on type III immunity.

**Figure 4.**
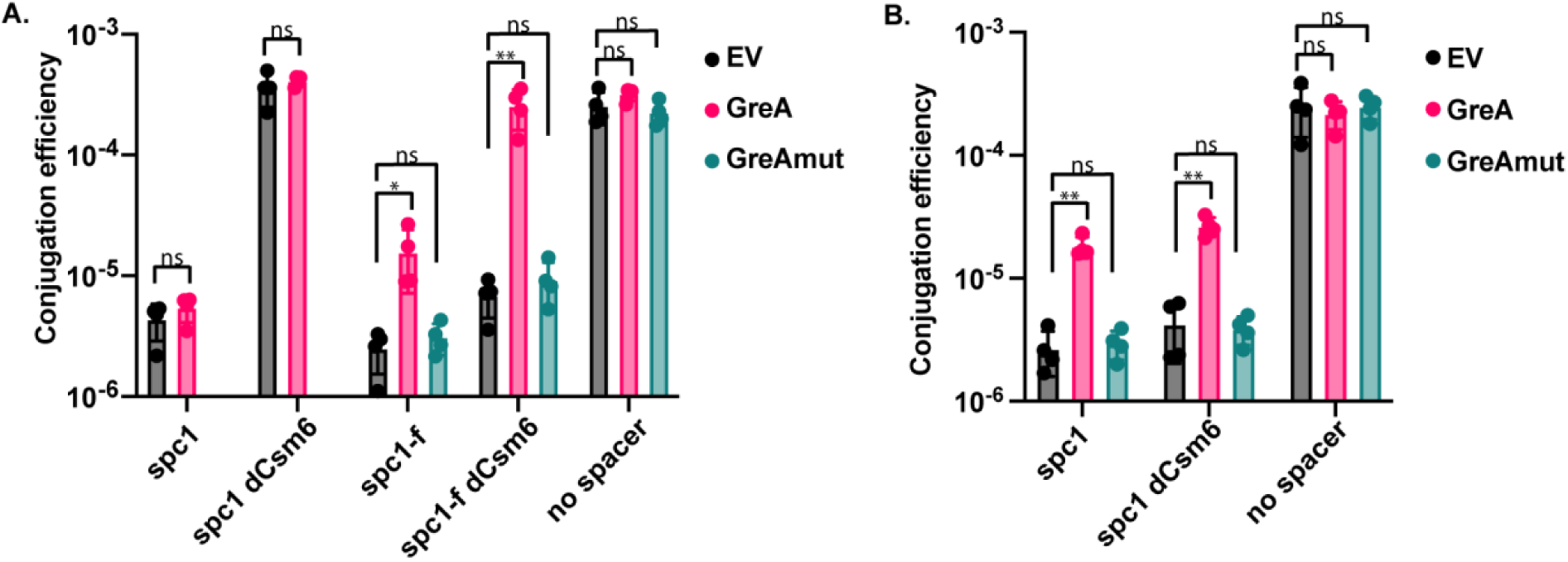
Anti-backtracking inhibits type III-A CRISPR anti-plasmid defense. **A.** Conjugation of pG0400 into cells carrying wild-type (WT) or Csm6-deactivated type III-A targeting the *nes* anti-sense transcript (spc1), the *nes* sense transcript (spc1-f), or without a targeting spacer (no spacer). Cells are also carrying either the GreA or GreAmut overexpression plasmids, or the empty vector (EV). Conjugation efficiency is calculated by dividing the amount of transconjugants by the amount of total recipient cells. Mean ± SD of 4 replicates is reported. **B.** Same as **A**, but with pG0420 being conjugated. pG0420 contains a promoter that drives high expression of the *nes* anti-sense transcript. Statistics were performed with an unpaired, two-sided t-test. ns: p>0.05; *: p<0.05; **: p<0.01; ***: p<0.001.

To test this hypothesis, we tested a type III system that targets the sense transcript of *nes* using a reverse spacer 1 (spc1-f). Under these conditions, GreA overexpression increased conjugation efficiency by approximately ten-fold, indicating a substantial reduction in anti-plasmid defense **(Figure 4A)**. We further examined pG0420, a derivative of pG0400 in which the antisense *nes* transcript is placed under the control of a strong promoter. In this background, GreA overexpression again increased conjugation efficiency by approximately 10-fold **(Figure 4B)**, demonstrating that GreA-mediated inhibition becomes apparent when target transcription is elevated.

We next investigated whether this phenotype was specifically associated with Cas10-mediated defense. Conjugation assays performed with a dCsm6 mutant, in which immunity relies exclusively on the Cas10 HD nuclease, revealed an even stronger effect: GreA overexpression essentially abolished anti-plasmid immunity, restoring pG0400 transfer to levels approaching those observed in the absence of CRISPR targeting (**Figure 4A)**. Mirroring the results with phage, overexpression of the catalytically inactive GreA mutant had no detectable effect on conjugation efficiency in either CRISPR-positive or CRISPR-negative strains (**Figure 4A and 4B)**. These findings demonstrate that inhibition of anti-plasmid immunity requires the anti-backtracking activity of GreA.

Together, these results extend our observations from phage infection to plasmid defense and demonstrate that RNA polymerase backtracking is also important for efficient type III immunity against conjugative elements. Notably, the inhibitory effect of GreA depended strongly on target transcription levels. GreA had little effect on immunity against the weakly transcribed native antisense *nes* target but substantially impaired defense when target transcription was increased or when immunity relied primarily on Cas10-mediated DNA degradation. One possible explanation is that RNAP backtracking contributes less to DNA targeting at weakly expressed loci. Alternatively, GreA may be recruited inefficiently to lowly transcribed genes, consistent with observations in *Streptococcus pneumoniae* showing that GreA has minimal effects on weakly expressed transcripts^35^. Together, these findings indicate that anti-backtracking activity selectively inhibits Cas10-dependent anti-plasmid immunity and that this effect is most pronounced at highly transcribed targets.

### Anti-backtracking of RNAP reduces DNA degradation caused by type III immunity

We next sought to directly visualize the impact of GreA on type III-mediated DNA degradation inside cells. In contrast to other CRISPR systems, DNA-targeting intermediates have not been isolated during type III immunity because target DNA is typically eliminated within minutes of target induction^12,10^. We used a *S. aureus* strain carrying an inducible target (*gp43*) sequence integrated into the chromosome (RN4220target)^36^. This strain was transformed with pCRISPR variants encoding *gp43* spacers complementary to the chromosomal target (**Figure 5A)**. Upon addition of anhydrotetracycline (aTc), transcription of the target sequence is induced, triggering type III CRISPR immunity against the host chromosome.

**Figure 5.**
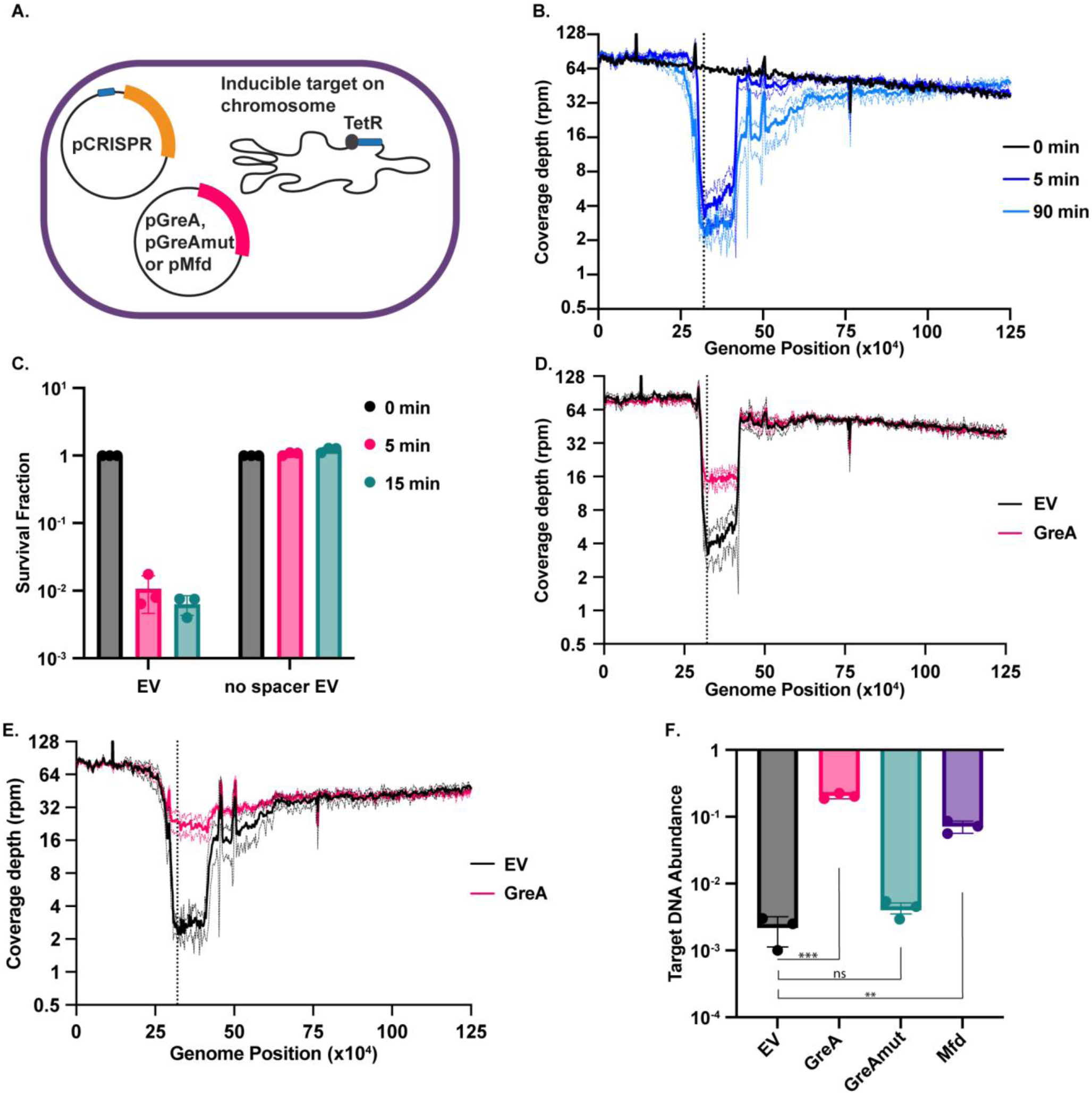
GreA overexpression reduces type III-A CRISPR-mediated DNA degradation. **A.** Schematic of the RN4220_Target_ strain. The bacterial chromosome contains an anhydrous tetracycline (aTc)-inducible promoter driving expression of the *gp43* protospacer of ΦNM1γ6. **B.** Coverage depth in reads per million (rpm) of RN4220_Target_ containing type III-A CRISPR with a *gp43* spacer before addition of aTc (0 min), and 5 minutes and 90 minutes after addition of aTc. Shown are the first 125 kB of the RN4220 genome, with position 0 is set to the origin of replication. Mean ± SD of 3 replicates is reported. Solid lines reflect the mean value while dotted lines show the ± SD. The black dotted vertical line shows the position of the inducible target. **C.** Survival fraction of RN4220_Target_ containing type III-A CRISPR with a *gp43* spacer or no spacer 5 and 15 minutes after aTc addition. Survival fraction is calculated by dividing the amount of CFU at the different time points by the amount of CFU before adding aTc (0 min). Mean ± SD of 3 replicates is reported. **D.** Same as **B** but with cells containing the GreA overexpression plasmid or the empty vector (EV) after 5 minutes of aTc addition. **E.** Same as (D) but 90 minutes after aTc addition. **F.** qPCR of the chromosomal *gp43* protospacer in RN4220_Target_ with a type III-A CRISPR with a *gp43* spacer 15 minutes after aTc addition. The cells also contained a GreA, GreAmut, or Mfd overexpression plasmid, or the empty vector (EV). Mean ± SD of 3 replicates is shown. Statistics were performed with an unpaired, two-sided t-test. ns: p>0.05; *: p<0.05; **: p<0.01; ***: p<0.001.

Following induction of the target transcript, we performed deep sequencing of the RN4220target genome at successive time points to monitor DNA degradation. At time 0, sequence coverage across the genomic region surrounding the target site was uniform, averaging approximately 100 reads per million (rpm). Strikingly, after only 5 min of induction, read coverage was markedly reduced across a ∼150 kb region centered on the target site, indicating rapid loss of chromosomal DNA from this locus (**Figure 5B**). By 90 min, the affected region had expanded to approximately 500 kb. Notably, DNA loss was highly asymmetric with respect to the target site. Read coverage recovered upstream of the target locus, whereas downstream coverage displayed a distinct stepwise pattern. The positions of these transitions corresponded closely to Chi sites in the *S. aureus* genome (5′-GAAGCGG-3′^37^) (**Figure S6A)**, which are known to attenuate the exonuclease activity of AddAB (also called RexAB in *S. aureus*)^38^. These observations suggest that AddAB contributes to double-stranded DNA degradation inside cells. Consistent with this model, the genomic region upstream of the target site contains a substantially higher density of Chi sites than the downstream region, providing a likely explanation for the asymmetric pattern of DNA loss. The extensive genomic degradation was accompanied by a loss of cell viability: RN4220target cells undergoing self-targeting exhibited approximately 99% killing within 5 min of target induction (**Figure 5C**), highlighting the rapid and destructive nature of type III-mediated chromosome targeting.

We next examined the contribution of each type III activity to chromosomal DNA degradation. Inactivation of the HD domain (dHD) completely abolished genome degradation, indicating that Cas10 nuclease activity is required for this process (**Figure S6B)**. Nevertheless, because Csm6 remained functional, RN4220target cells carrying the dHD mutant still exhibited a reduction in viability following target induction, although killing was less pronounced than in the wild-type system (**Figure S6D)**. In contrast, inactivation of Csm6 (dCsm6) had little effect on genome degradation, which closely resembled that observed in the wild-type and was accompanied by a comparable loss of viability (**Figure S6C and S6D)**. Together, these results demonstrate that the Cas10 HD domain is both necessary and sufficient for type III-mediated DNA degradation *in vivo*, whereas Csm6 contributes to cell killing through a DNA degradation-independent mechanism.

Using this system, we asked whether overexpression of factors that suppress RNAP backtracking influences the extent of type III-mediated DNA degradation. RNA-seq on RN4220target in the presence or absence of GreA overexpression showed that the transcript level of the region surrounding the inducible target were not significantly different between the conditions, suggesting that GreA expression did not affect target transcription (**Figure S7A)**. During type III self-targeting, overexpression of GreA markedly reduced genomic DNA degradation in RN4220target cells relative to the empty-vector control (**Figure 5D, 5E, and 5F**). Consistent with this reduction in DNA damage, self-targeting cells expressing GreA exhibited significantly improved survival compared to cells carrying the empty-vector control (**Figure S7B)**. Similarly, overexpression of Mfd reduced DNA degradation, as measured by qPCR (**Figure 5E)**, and correspondingly increased bacterial survival (**Figure S7B**). In contrast, expression of a catalytically inactive GreAmut failed to alter either DNA degradation or cell viability (**Figure 5F and S7B)**, both of which remained indistinguishable from the empty-vector control. Similarly to the phage system, RNaseHIII overexpression had no detectable effect on either type III-mediated DNA degradation or cell survival in the RN4220target strain (**Figure S7C and S7D)**. Together, these results indicate that suppression of RNAP backtracking, rather than resolution of R-loops, limits type III CRISPR DNA degradation and the associated loss of viability.

### Phage genomes encode for GreA-like factors

Mobile genetic elements frequently encode RNAP-binding proteins that reprogram host transcription. For example, both the conjugative F plasmid and bacteriophage λ of *Escherichia coli* encode TraR, a transcription factor with structural similarity to GreA and other RNAP-binding regulators such as DksA^39^. Like GreA, TraR consists of a globular C-terminal domain connected by a flexible linker to an N-terminal α-helix, although the corresponding N-terminal region of GreA forms an elongated coiled-coil structure40. In both plasmid F^41^ and phage λ^39^, TraR binds RNAP and modulates transcription during their respective life cycles. These observations prompted us to investigate whether bacteriophages also encode GreA homologs.

To identify phage-encoded GreA proteins, we searched the IMG/VR v4 database of high-quality viral genomes^42^ using three HMM profiles corresponding to the conserved domains of GreA. Candidate sequences were retained only if they were either located on circular viral genomes or flanked within 10 kb on both sides by genes encoding viral proteins, minimizing the likelihood of host-derived or non-viral sequence contamination. This analysis identified 122 phage genomes encoding at least one GreA-related domain. Of these, 76 encoded GreA-like proteins ranging from 70 to 260 amino acids in length (compared to 158 amino acids for *E. coli* GreA) and were used to construct a phylogenetic tree (**Figure S8**).

Among these 76 candidates, 54 retained the hallmark catalytic motif of GreA, consisting of an aspartate and glutamate separated by two residues (DXXE) within a predicted N-terminal coiled-coil domain. Sequence alignments of the three highest-scoring candidates, based on similarity to GreA from *E. coli* and *S. aureus*, are shown in **Figure 6A**. Consistent with their sequence conservation, AlphaFold3-predicted structures revealed strong structural similarity to bacterial GreA proteins, with root-mean-square deviations (RMSDs) ranging from 0.844 to 0.876 Å relative to the *E. coli* GreA structure (**Figure 6B**). Together, these results suggest that GreA-like transcription factors are widespread among bacteriophages and retain the key structural features required for GreA function.

**Figure 6.**
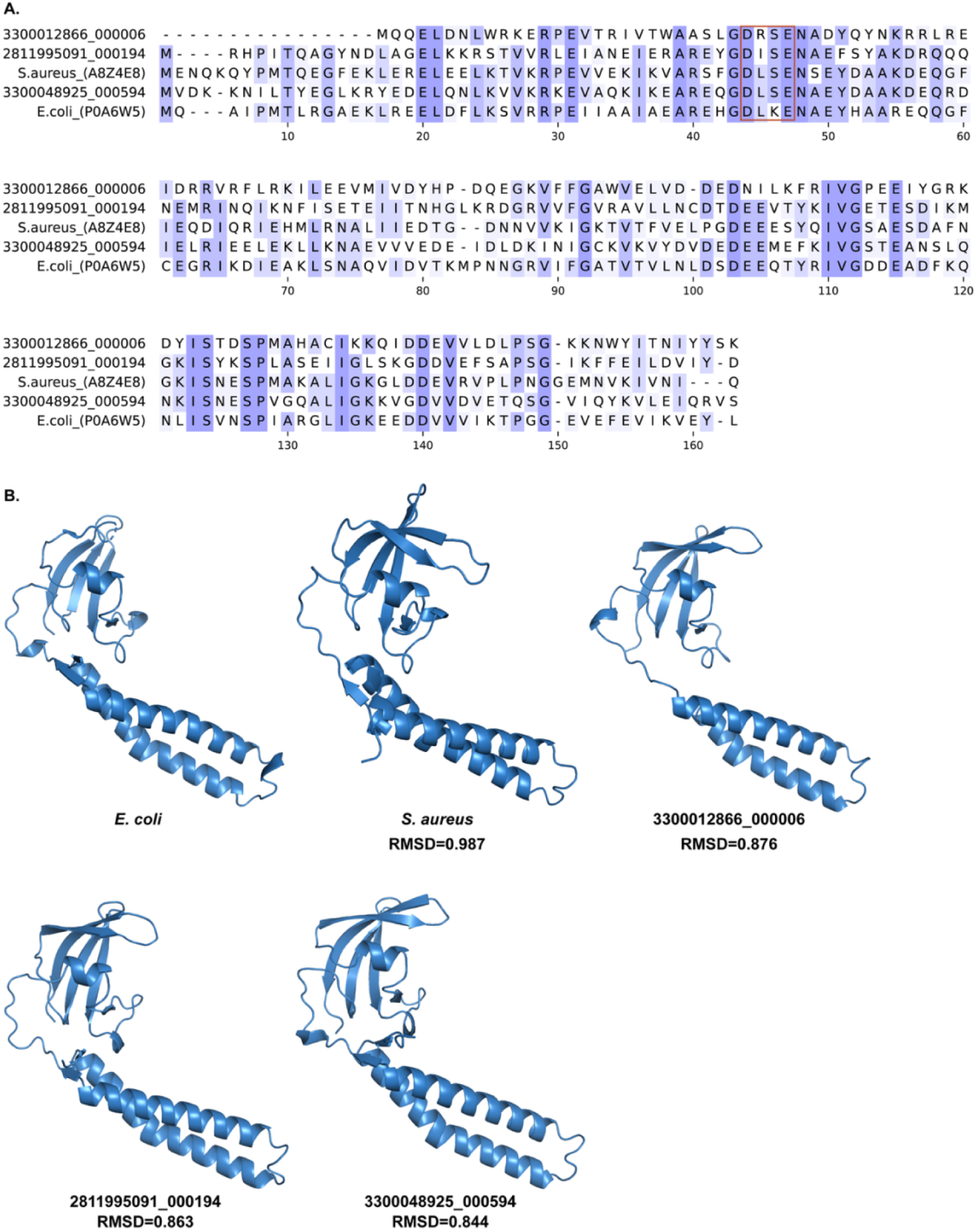
Phage genomes from IMG/VR database encode for GreA*-*like sequences. **A.** Amino acid sequence alignment of *Escherichia coli* (UniProt: P0A6W5) and *S. aureus* (UniProt: A8Z4E8) GreA, as well as the top three phage-encoded GreA by sequence similarity (IMG/VR identifier is shown). The red rectangle contains the conserve aspartic and glutamic acid required for the anti-backtracking function of GreA. **B.** Structure of *E. coli* GreA (UniProt: P0A6W5, structure ID 1GRJ), as well as the AlphaFold3 predicted structure of *S. aureus* GreA (UniProt: A8Z4E8) and of the three phage-encoded GreA-like sequences from **A**. RMSD were obtain aligning each predicted structure to *E. coli* GreA.

Phages can encode for anti-CRISPR proteins that allow them to evade CRISPR immunity^43^. Studies have shown that anti-CRISPR proteins are usually small and can be co-opted from bacterial host encoded genes^44^. We therefore asked whether the host bacteria of the 76 phages encoding with GreA-like sequences carried any type III CRISPR-Cas systems. We used blastp^45^ to searched for bacterial homologs of all 76 phage-encoded GreA-like proteins to identify their closest cellular counterparts, reasoning that these genes may have been acquired from bacterial hosts through horizontal gene transfer. For each candidate, the taxonomic assignment of the predicted host reported in IMG/VR matched the taxonomy of the top BLASTP hit, supporting a host-associated origin for these proteins.

To further explore a potential relationship between phage-encoded GreA-like proteins and CRISPR-Cas immunity, we surveyed publicly available genomes from NCBI representing the bacterial genera predicted to host phages carrying GreA-like sequences and determined the CRISPR-Cas systems present within each genus. Of the predicted host genera, 13 contained type III-A CRISPR-Cas systems (**Figure S9**). Although correlative, these findings raise the intriguing possibility that some phage-encoded GreA-like proteins function as anti-type III defense factors. Given the established role of GreA in suppressing type III immunity, phages may exploit GreA-like proteins to interfere with host CRISPR-Cas activity during infection. However, experimental validation will be required to determine whether these proteins directly antagonize type III CRISPR-Cas systems.

## Discussion

In this study, we show that anti-backtracking factors, such as GreA and Mfd, can inhibit type III-A CRISPR-Cas immunity. Overexpression of either factor reduced Cas10-Csm HD domain-mediated DNA targeting, thereby permitting replication of diverse phages and plasmids. To our knowledge, this is the first demonstration that RNA polymerase backtracking directly influences type III CRISPR immunity, revealing a previously unrecognized mechanistic link between transcription dynamics and type III immune function and providing new insight into how both bacteria and mobile genetic elements may regulate the activity of this widespread defense system.

A defining feature of type III CRISPR-Cas systems is their dual mode of immunity, which combines RNA degradation with DNA targeting. Previous studies have identified both bacterial and phage-encoded factors that modulate type III immune responses; notably, however, most of these factors act on the RNA-targeting arm of the pathway. For example, the bacterial RNases RNaseR contribute to type III immunity against plasmids by cooperating with Csm6 to degrade target transcripts^46^. Likewise, several phages encode factors that degrade or prevent synthesis of cyclic oligoadenylate (cOA) molecules synthesized by the Cas10 PALM domain, thereby preventing activation of the Csm6 RNase and functioning as anti-CRISPR proteins^47^. One notable exception is an anti-CRISPR system described by the Hatoum-Aslan laboratory that directly binds to Cas10-Csm and disables the entire effector complex^48^. Our findings reveal a distinct mechanism of regulation. By controlling RNA polymerase backtracking, GreA and Mfd selectively modulate the DNase activity of the type III system: overexpression of either factor impaired immunity mediated by the Cas10 HD domain while leaving RNA-targeting or Cas9-mediated immunity unaffected. These results identify RNAP backtracking as a mechanism for specifically regulating type III DNA targeting and suggest that transcription dynamics can tune the relative contributions of DNA targeting in type III immunity. Indeed, we found GreA-like sequences encoded in phage genomes recovered from metagenomic data. Future work will determine whether these sequences are functional anti-backtracking factors and whether they can inhibit type III CRISPR immunity.

Our work offers a new model for how type III CRISPR-mediated DNA targeting occurs *in vivo*. Despite extensive biochemical and structural studies on of Cas10-Csm, it remains unclear how the complex accesses DNA during transcription. The exposed non-template strand within an actively transcribing RNA polymerase elongation complex is only ∼9 nucleotides long and is transiently available as the polymerase moves on average at a rate of 30–100 nucleotides per second^49,50^. Our data raise the possibility that backtracked RNA polymerases provide a more accessible substrate for Cas10-Csm DNA targeting; inhibiting backtracking thus limits DNA targeting. We therefore propose that RNAP backtracking facilitates type III DNase activity by increasing access to exposed ssDNA. Future studies will be needed to determine how backtracked transcription complexes promote DNA targeting and whether Cas10-Csm itself can induce or stabilize RNAP backtracking.

Our work also provides a mechanism by which type III degrades double-stranded DNA targets. Previous work has shown that type III can degrade plasmid and phage DNA within minutes of target induction, yet no degradation intermediates were detected. This contrasts with type I and type II CRISPR systems, where sequencing-based approaches have identified well-defined cleavage sites and degradation products. Moreover, conflicting evidence has raised questions about whether DNA depletion results directly from Cas10 ssDNase activity or indirectly from Csm6-mediated growth arrest. Using the chromosomal self-targeting system, we demonstrated that type III-A CRISPR immunity drives progressive DNA degradation surrounding the target site in a Cas10 HD domain-dependent manner. The stepwise pattern of DNA loss at Chi sites suggests the involvement of the AddAB helicase-nuclease complex. AddAB recognize double-stranded DNA breaks and processively degrade DNA until they encounter Chi sites. We therefore propose a model in which Cas10-Csm initially generates a single-stranded DNA lesion that is subsequently converted into a double-stranded break, thereby recruiting AddAB to degrade both strands of the target genome. These findings could assist in the development of novel DNA editing tools that leverage type III transcription-dependent DNA degradation and anti-backtracking mediated inhibition.

We acknowledge that alternative interpretations of our results and model are possible. Because GreA functions broadly as an RNAP fidelity factor, its overexpression could, in principle, influence type III immunity through mechanisms independent of Cas10-Csm. However, our transcriptomic analyses argue against a global effect on host gene expression. We also consider it unlikely that GreA inhibits AddAB-mediated DNA degradation, as Sivaramakrishnan et al. showed that GreA promotes – not inhibits – the analogous RecBCD-dependent degradation pathway in *E. coli*^51^. Another possibility is that reduced RNAP backtracking enhances repair of the targeted DNA sequence, yet this explanation is inconsistent with previous findings that GreA impairs DNA break repair^51^. Moreover, GreA overexpression neither diminished Cas9-mediated targeting nor increased the frequency of escaper mutants during type III or type II targeting. Together, these observations support the conclusion that the inhibitory effect of GreA is specific to the Cas10-Csm complex rather than a consequence of broader effects on transcription or DNA repair.

## Acknowledgements

We thank the University of Wisconsin – Madison Biotechnology Center’s Gene Expression Core (Research Resource Identifier – RRID:SCR_017757) for RNA library preparation and sequencing. We gratefully acknowledge Luciano Marraffini for providing bacterial and phage stocks; Robert Landick, Jade Wang, Jason Peters, and David Hershey for helpful experimental advice; and all members of the Mo group for their feedback and support. AJHV and JCK are supported by individual NSF Graduate Research Fellowships; NG is supported by the Hilldale Undergraduate Research Fellowship; CYM is supported by startup funds from the Department of Bacteriology at UW-Madison and the Margeret Q. Landenberger Foundation; KA is supported by a National Institute of General Medical Sciences of the National Institutes of Health award (R35GM143024).

## Methods

### Bacterial strains and growth conditions

*Staphylococcus aureus* (strains RN4220 or TB4) were culture in Brain Heart Infusion (BHI) medium. When appropriate, the medium was supplemented with chloramphenicol (10 μg/mL) to select for plasmids encoding CRISPR systems, with erythromycin (10 μg/mL) to select for the overexpression plasmid (and the empty vector), and with mupirocin (5 μg/mL) to select for the pG0400 or pG0420 conjugative plasmid. Strains were grown at 37°C with shaking at 250rpm.

For plated cultures, BHI agar was supplemented when appropriate with with chloramphenicol (10 μg/mL) to select for plasmids encoding CRISPR systems, with erythromycin (10 μg/mL) to select for the overexpression plasmid (and the empty vector), and with mupirocin (5 μg/mL) to select for the pG0400 or pG0420 conjugative plasmid. All plates were incubated at 37°C.

### Plasmid cloning

pCYM014 was constructed using Gibson assembly by introducing the pHyperSpank promoter into the shuttle vector pCN057. In brief, the pHyperSpank promoter along with a multiple cloning site and lacI were amplified with PCR with primers CYM-Pm075 and CYM-Pm076, while the pCN057 was amplified with primers CYM-Pm073 and CYM-Pm074. The two DNA fragments were joined in a two-fragment Gibson reaction following the protocol by the NEBuilder Hifi DNA Assembly (NEB: E2621).

The GreA, Mfd, and RNaseHIII overexpression plasmids (pCYM108, pCYM107, and pAH09 respectively) were built by amplifying those genes and flanking regions from RN4220 using primers pairs CYM-Pm482/CYM-Pm483 (for *greA*), CYM-Pm480/CYM-Pm481 (for *mfd*), and oAH035/oAH036 (for *rnhC*), respectively; and joined by Gibson assembly with a pCYM014 backbone that was restriction-digested with PstI-HF (NEB: R3140) and SpeI-HF (NEB: R3133) using the NEBuilder Hifi DNA Assembly. The GreAmut overexpression plasmid was constructed using Gibson assembly by joining a gBlock (see Supplementary Sequence 1) ordered from Integrated DNA Technologies (IDT) containing the mutant *greA* sequence allele as well as Gibson junctions into a pCYM014 backbone that was restriction-digested with PstI-HF and SpeI-HF using the NEBuilder Hifi DNA Assembly. pCYM130 was constructed by digesting pGG-BsaI-R with BsaI-HF (NEB: R3733) and ligating this vector with T4 ligase (NEB: M0202) to annealed complementary spacer oligos (spc59F and spc59R).

### Preparation of electrocompetent *S. aureus*

Overnight cultures of *S. aureus* were diluted 1:100 in 25 mL of BHI medium and incubated at 37°C with shaking at 250 rpm until an OD600 of ∼0.7 was reached. The culture was washed with water 4 times, then a final wash with 10% glycerol was performed. The cells were then resuspended in 250 μL of 10% glycerol.

### Purification of plasmid DNA

Plasmid DNA was purified from 5mL of overnight culture of *S. aureus.* The culture was pelleted and resuspended in 250 μL of buffer P1from QIAprep Spin Miniprep Kit (Qiagen, catalog number 27104) and 50 μL of lysostaphin (1 mg/mL). After a one hour incubation period at 37°C, the bacterial lysate was treated with the reagents from QIAprep Spin Miniprep Kit. Plasmid DNA was eluted using 35 μL of water.

### Transformation of *S. aureus*

50 μL of electrocompetent cells were mixed with 3 μL of plasmid DNA. The cells were electroporated using standard conditions for staphyloccoci. After electroporation, 1 mL of BHI was added to the cells and incubated at 37°C with shaking at 250 rpm for an hour before plating in BHI agar plates with the appropriate antibiotic for selection of transformants.

### Phage plaquing assay

200μL of overnight cultures were mixed with 5mL of BHI top agar (0.5x BHI agar) and pour over BHI plates. After the top agar solidified, 3 μL of different concentrations of phage were spotted onto the top agar. Plates were incubated at 37°C for ∼16 hours.

### Average Burst Size Assay

Measurement of average burst size was performed as described previously ^10^. In brief, *S. aureus* cells were grown in BHI with 5mM CaCl2 and appropriate antibiotics to an OD600 of 0.3. At this point IPTG (final concentration of 1 mM) was added and infected by ΦNM1γ6 at MOI = 0.1 or with Φ12 at MOI=1 for 5 min. Cells were immediately washed in BHI twice at 4°C and re-suspended in equal initial volume. An aliquot of cells were spotted on BHI top agar plates with RN4220. The rest of the cells were incubated at 37°C for another 75 min before an aliquot of cells were spotted on RN4220. Plates were incubated at 37°C for ∼16 hours before plaques were counted. Average burst size was calculated as the ratio of plaques formed at 80 min to plaques formed at 5 min for each strain of interest.

### Bacterial growth curve during phage infection

Overnight cultures of *S. aureus* were diluted 1:100 into BHI with 5 mM CaCl2 and with the appropiate antibiotics. After the cultures reached an OD600 of 0.2-0.3, aliquotes of the cultures were transferred into 96-well plates, IPTG was added for a final concentration of 1 mM and phage was added at the indicated MOI. The 96-well plate was put into a shaking plate reader at 37°C and OD600 was measured every 20 minutes.

### Phage Escaper Assay

Phage escapers were quantified as described previously^34^. In brief, overnight cultures of *S. aureus* with CRISPR and overexpression pasmids were diluted 1:100 into 3 mL of BHI with 5 mM CaCl2, when the culture reached an OD600 of 0.3, phage was added at an MOI=20. Immediately after, an aliquote of the cells were spotted on BHI top agar plates containing cells with CRISPR plasmids (for quantification of phage escapers) and on BHI top agar plates with RN4220 (for quantification of total phage). This was repeated after 4 hours of incubation. Immedietly after, the cultures were pelleted and the supernatant containing the phages was isolated. Exponentially growing *S. aureus* with CRISPR plasmids and overexpression plasmids were added to the supernatant to a final OD600 of 0.3. These cultures were incubated for 4 more hours (8 hour post infection), and both phage escapers and the total amount of phage was quantified.

### Conjugation assays

Overnight cultures of donor and recipient strains were diluted 1:100 and grown at 37°C with shaking at 250 rpm. When the cultures reached an OD600 of ∼0.8, 500 μL of donor and 500 μL of recipient were mixed and washed with the fresh BHI. The culture was resuspended on 100 μL of fresh BHI and plated on top of a 0.45 μm filter paper on BHI agar plates. The plates were incubated at 37°C for 24 hours. The filter paper containing the mixed culture was then resuspended in 5 mL of BHI, serially diluted for CFU counting, and plated on BHI agar plates with chloramphenicol and erythromycin for total recipient count, and on chloramphenicol and muporocin for transconjugants count. The conjugation efficiency was calculated by dividing the amount of transconjugations by the amounts of total recipients.

### Total DNA extraction

Phage and bacterial DNA was purified using phenol-chloroform extraction. To lyse *S. aureus* 10 mL of exponentially growing cells were pelleted and resuspended in 500 μL of P1 Buffer (from QIAprep Spin Miniprep Kit, Qiagen catalog number 27104) and 50μL of lysostaphin (1 mg/mL). DNA was resuspended in 200 μL of water.

### DNA-sequencing of RN4220Target and coverage depth analysis

Overnight cultures of RN4220Target strain were diluted 1:100 in 50 mL of BHI and incubated at 37°C with shacking until they reached an OD600 of 0.5 before anhydrous tetracycline (aTc) was added for a final concentration of 125 ng/mL. 10mL Aliquots for DNA extraction were taken just before adding aTc, and 5 minutes, and 90 minutes after aTc addition. DNA samples were sent to SeqCenter for Illumina sequencing. Genomic assembly was performed on Geneious Primer (version 2026.0.2) using RN4220Target as a reference genome. We then extracted a table from Geneious Primer containing the read coverage for each genomic position. We divided the reads of each position by the total number of reads and then multiplied by a million to calculate reads per million (rpm). The rpm was binned for every 2,000 positions.

### qPCR

The qPCR assays were performed using the SsoAdvanced universal SYBR Green supermix (Biorad, 172-5270∼5275), and performed according to their instructions. 75 ng of DNA were used in each reaction. Reactions were done on Biorad CFX96 real-time PCR instrument. For the qPCR during phage infection, overnight cultures were diluted 1:100 in 50 mL and grown at 37°C with shacking until they reached an OD600 of 0.3. Phage was added at an MOI of 5. 10mL aliquots of culture were taken 15 minutes, 45 minutes, 1.5 hours, and 3 hours after addition of phage for total DNA extraction. The 2^-ΔΔCq^ method was used to analyze the results. The bacterial host gene rho was used as a reference gene, the data was normalized to the average of the Cq values of the 3 replicates of the phage gene *gp14* of the empty vector condition at 15 minutes after phage addition.

For the qPCR of RN4220Target, overnight cultures were diluted 1:100 in 50 mL and grown at 37°C with shacking until they reached an OD600 of 0.5. 10mL aliquots were taken at this point for total DNA extraction. Then, anhydrous tetracycline (aTc) was added for a final concentration of 125 ng/mL. 15 minutes later, 10mL aliquots were taken for total DNA extraction. The 2^-ΔΔCq^ method was used to analyze the results.The rho gene was used as a reference, and the data was normalized to the average of the Cq values of the 3 replicates of the target locus before adding aTc.

### Bacterial Survival Curves

Overnight cultures of RN4220Target were diluted 1:100 and grown at 37°C with shaking at 250 rpm. When cultures reached and OD600 of 0.5, an aliquot was taken and diluted for CFU counting. Immediately after, anhydrous tetracycline (aTc) was added for a final concentration of 125 ng/mL. 5 minutes and 15 minutes after aTc addition, aliquots were taken a diluted for CFU counting.

### Total RNA extraction and RNA-sequencing

Overnight cultures of RN4220Target strain were diluted 1:100 in BHI and incubated at 37°C with shacking until they reached an OD600 of 0.7 before anhydrous tetracycline (aTc) was added for a final concentration of 125 ng/mL. After 5 minutes of aTc addition, 500 µL of culture were transferred into a microcentrifuge containing 1 mL of RNAprotect Bacteria reagent (QIAGEN: 76506) was added. The mixture was vortexed for 5 seconds and incubated at room temperature for 5 minutes. The tubes were centrifuged for 10 mins at 5000 × *g*, and the supernatant discarded. For the RNA extraction, bacterial pellets were resuspended in 80 µL of phosphate buffered saline and 10 µL of lysostaphin solution (1 mg/mL stock). The suspensions were incubated at 37 °C, with shaking, until the solution looked clear (∼30 minutes). 10 µL of 10% sarkosyl was then added and the tube mixed, after which 300 µL of TRIzol reagent (Invitrogen: 15596026) was added. RNA purification was performed following the protocol from the Direct-zol RNA Miniprep Plus kit (ZYMO Research: R2050). DNase I treatment was performed as recommended, and the samples were eluted in 50 µL of DNase/RNase-free water. Purified RNA was prepared and sequenced on an Illumina sequencing platform at the UW-Madison Gene Expression Center.

### Differential gene expression analysis

RNA was collect from RN4220Target containing a CRISPR plasmid without a targeting spacer, with the GreA overexpression plasmid or the empty vector control (EV) to assess the impact of GreA on the strains transcriptome. Cleaned RNA reads were mapped to the RN4220Target reference genome using bowtie2^52^ v2.5.4. FeatureCount^53^ v2.0.8 was used to generate a read count matrix, which was analyzed with DESeq2^54^ v1.44.0. Figures were generated using the packages tidyverse (version 2.0.0) and EnhancedVolcanoPlots (version 1.22.0).

### Search of GreA-like sequences in phage genomes

Viral genomes were obtained from the IMG/VR v4 database42 and filtered to include only “high-quality” sequences, as determined by CheckV^55^, with a minimum genome length of 5,000 bp enforced. Open reading frames in the filtered set of genomes were predicted with pyrodigal-gv, an extension of pyrodigal^56^, the Python bindings and interface to Prodigal^57^, that adds the virus gene models introduced with geNomad^58^. Only genomes encoding at least four proteins were included in the analysis described below.

To annotate the functions of proteins encoded by viral genomes, we used a pre-release version of CheckAMG^59^ (October 2024). All virus-encoded proteins were searched with PyHMMER v0.10.10^60^, a Python library binding to HMMER^61^, against profile HMMs from PHROG v4^62^ (release 2022-01-17), KEGG KOfam^63^ (March 2019 release) and Pfam-A v35.0^64^. Each database was searched separately, and a minimum alignment coverage of 50% of the viral protein sequence was enforced. When a protein had more than one hit to profile HMMs within the same database, only the hit with the highest bit score was retained.

To search specifically for GreA-like proteins, we used a set of three profile HMMs from the Pfam database^64^: PF01272.19 (Transcription elongation factor, GreA/GreB, C-term), PF03449.15 (Transcription elongation factor, N-terminal) and PF20698.2 (PIN domain associated with the TPR-GreAB-C-PIN system), where PF01272.19 and PF03449.15 are Pfam 32.0-era builds. These three profile HMMs were searched as a custom set in the CheckAMG pipeline, separate from the annotation databases described above. To remove partial alignments and weak hits, a protein was retained as a GreA-like protein when its alignment to a reference profile HMM covered at least 50% of the profile HMM, had an E-value of at most 1e^-5^, and reached at least 25% amino acid identity.

The GreA-like proteins were then conservatively filtered to minimize the possibility that they originated from host genome contamination. This was achieved by assigning V-scores and VL-scores to the annotated proteins of the viral genomes ^65^, since these metrics indicate whether a protein is typically encoded by viruses or by non-viral sources. Using the best profile HMM hit of each viral protein, we required each GreA-like protein to be flanked on both sides, within 10 kb, by a protein whose profile HMM annotation carried a V-score of 10, which signifies a strong viral association ^65^. GreA-like sequences encoded on a circular viral genome, identified by CheckAMG from terminal repeats, were also accepted. Additionally, the local genomic region encoding the GreA-like protein (a 10 kb window around the gene) was required to have an average VL-score of at least 3.15, which indicates that the region as a whole is of viral origin ^65^. GreA-like proteins passing all filters were retained for further analysis.

### AlphaFold3 modeling

Putative GreA-like proteins were predicted using AlphaFold3^66^ on the Center for High Throughput Computing’s cluster in March 2025. Input protein sequences were provided in fasta format and then converted to the recommended JSON format with dialect Alphafold3^66^, version 1, and model seed 1 for all sequences. 5 models were generated for each protein sequence, and ranked_scores were obtained. The best model was picked for each prediction (rank 0). Protein structures were visualized in PyMol (PyMOL, The PyMOL Molecular Graphics System, Version 3.0 Schrödinger, LLC) and RMSD values were calculated using the “align” function.

### Predicted host analysis

We used blastp^45^ on all 76 greA-like protein sequences against the NCBI clustered_nr database, and reported the best match and accession ID. The taxonomy of the top hit was used to report the taxonomy of the putative bacterial host. In addition, since the predicted greA-like sequences were from the IMG VR database^42^, we pulled predicted host information from IMG/VR (https://img.jgi.doe.gov/cgi-bin/vr/main.cgi) using the function Search UVIG ID, and provided the comma-separated list of IMG UViG ID (for example: IMGVR_UViG_XXXXX_XXXXXX). The table was exported **(Supplementary File 2).** 32 out of 76 UViG IDs had a predicted host taxonomy as per IMG/VR. In any case, the reported taxonomy of the 32/76 GreA-like sequences matched the BLASTp top-hit taxonomy.

### GreA phylogenetic tree

We created a greA gene phylogeny using the 76 phage greA protein sequences, along with two bacterial greA references (*E. coli* Uniprot P0A6W5 and Staphylococcus aureus Uniprot A8Z4E8), for a total of 78 sequences. The alignment was created using MAFFT v7.490^67^ (Automatic algorithm, Blosum62 Scoring matrix, gap open penalty 1.53, offset value 0.123).

IQ-TREE version 3.1.3^68^ was used to generate a greA gene phylogeny, and to pick the best model as identified by ModelFinder^69^ using the following command: iqtree3 -s [multiple sequence alignment file] greA-76.fasta -alrt 1000 -m MFP -B 1000 -T AUTO. The best-fit model according to model metrics, according to BIC (Bayesian Information Criterion), was LG+R3. The random seed used was 224025. The consensus tree containing the bootstrap values was visualized using the program iTOL.

### Identification and classification of CRISPR systems in bacterial genomes

We searched for genomes belonging to the taxonomic groups of the predicted host taxonomy on the Genome Taxonomy Database (GTDB). From these tables, each genomes had both a GTDB classification and a NCBI name, along with the exact genome accession ID from the NCBI GenBank database. We then removed duplicate accession numbers, excluded any genomes that failed the quality check, and kept accession numbers that were GTDB representative genomes or NCBI type species belonging to the genus of interest – resulting in 16294 accession IDs. (**Supplementary File 3**). 16294 genome records were found and downloaded using NCBI’s datasets tool. We ran cctyper^70^ on the 16294 genomes on the Center for High Throughput Computing HTC cluster. The output files ending with “cas_operons.tab” were combined, analyzed, and visualized in R.

## References

1. Barrangou, R., Fremaux, C., Deveau, H., Richards, M., Boyaval, P., Moineau, S., Romero, D.A., and Horvath, P. (2007). CRISPR Provides Acquired Resistance Against Viruses in Prokaryotes. Science 315, 1709–1712. 10.1126/science.1138140.

2. Marraffini, L.A. (2015). CRISPR-Cas immunity in prokaryotes. Nature 526, 55–61. 10.1038/nature15386.

3. Makarova, K.S., Aravind, L., Wolf, Y.I., and Koonin, E.V. (2011). Unification of Cas protein families and a simple scenario for the origin and evolution of CRISPR-Cas systems. Biol. Direct 6, 38. 10.1186/1745-6150-6-38.

4. Makarova, K.S., Shmakov, S.A., Wolf, Y.I., Mutz, P., Altae-Tran, H., Beisel, C.L., Brouns, S.J.J., Charpentier, E., Cheng, D., Doudna, J., et al. (2025). An updated evolutionary classification of CRISPR–Cas systems including rare variants. Nat. Microbiol. 10, 3346–3361. 10.1038/s41564-025-02180-8.

5. Tamulaitis, G., Kazlauskiene, M., Manakova, E., Venclovas, Č., Nwokeoji, A.O., Dickman, M.J., Horvath, P., and Siksnys, V. (2014). Programmable RNA Shredding by the Type III-A CRISPR-Cas System of Streptococcus thermophilus. Mol. Cell 56, 506–517. 10.1016/j.molcel.2014.09.027.

6. Kazlauskiene, M., Tamulaitis, G., Kostiuk, G., Venclovas, Č., and Siksnys, V. (2016). Spatiotemporal Control of Type III-A CRISPR-Cas Immunity: Coupling DNA Degradation with the Target RNA Recognition. Mol. Cell 62, 295–306. 10.1016/j.molcel.2016.03.024.

7. Niewoehner, O., Garcia-Doval, C., Rostøl, J.T., Berk, C., Schwede, F., Bigler, L., Hall, J., Marraffini, L.A., and Jinek, M. (2017). Type III CRISPR–Cas systems produce cyclic oligoadenylate second messengers. Nature 548, 543–548. 10.1038/nature23467.

8. Mo, C.Y., Mathai, J., Rostøl, J.T., Varble, A., Banh, D.V., and Marraffini, L.A. (2021). Type III-A CRISPR immunity promotes mutagenesis of staphylococci. Nature 592, 611–615. 10.1038/s41586-021-03440-3.

9. Jia, N., Jones, R., Yang, G., Ouerfelli, O., and Patel, D.J. (2019). CRISPR-Cas III-A Csm6 CARF Domain Is a Ring Nuclease Triggering Stepwise cA4 Cleavage with ApA>p Formation Terminating RNase Activity. Mol. Cell 75, 944–956.e6. 10.1016/j.molcel.2019.06.014.

10. Jiang, W., Samai, P., and Marraffini, L.A. (2016). Degradation of Phage Transcripts by CRISPR-Associated RNases Enables Type III CRISPR-Cas Immunity. Cell 164, 710–721. 10.1016/j.cell.2015.12.053.

11. Aviram, N., Shilton, A.K., Lyn, N.G., Reis, B.S., Brivanlou, A., and Marraffini, L.A. (2024). Cas10 relieves host growth arrest to facilitate spacer retention during type III-A CRISPR-Cas immunity. Cell Host Microbe 32, 2050–2062.e6. 10.1016/j.chom.2024.11.005.

12. Rostøl, J.T., and Marraffini, L.A. (2019). Non-specific degradation of transcripts promotes plasmid clearance during type III-A CRISPR–Cas immunity. Nat. Microbiol. 4, 656–662. 10.1038/s41564-018-0353-x.

13. Samai, P., Pyenson, N., Jiang, W., Goldberg, G.W., Hatoum-Aslan, A., and Marraffini, L.A. (2015). Co-transcriptional DNA and RNA Cleavage during Type III CRISPR-Cas Immunity. Cell 161, 1164–1174. 10.1016/j.cell.2015.04.027.

14. Liu, T.Y., Iavarone, A.T., and Doudna, J.A. (2017). Correction: RNA and DNA Targeting by a Reconstituted Thermus thermophilus Type III-A CRISPR-Cas System. PLOS ONE 12, e0175612. 10.1371/journal.pone.0175612.

15. Lin, J., Shen, Y., Ni, J., and She, Q. (2021). A type III-A CRISPR–Cas system mediates co-transcriptional DNA cleavage at the transcriptional bubbles in close proximity to active effectors. Nucleic Acids Res. 49, 7628–7643. 10.1093/nar/gkab590.

16. Liu, T.Y., Liu, J.-J., Aditham, A.J., Nogales, E., and Doudna, J.A. (2019). Target preference of Type III-A CRISPR-Cas complexes at the transcription bubble. Nat. Commun. 10, 3001. 10.1038/s41467-019-10780-2.

17. Landick, R. (2021). Transcriptional Pausing as a Mediator of Bacterial Gene Regulation. Annu. Rev. Microbiol. 75, 291–314. 10.1146/annurev-micro-051721-043826.

18. Agapov, A., Olina, A., and Kulbachinskiy, A. (2022). RNA polymerase pausing, stalling and bypass during transcription of damaged DNA: from molecular basis to functional consequences. Nucleic Acids Res. 50, 3018–3041. 10.1093/nar/gkac174.

19. James, K., Gamba, P., Cockell, S.J., and Zenkin, N. (2016). Misincorporation by RNA polymerase is a major source of transcription pausing *in vivo*. Nucleic Acids Res., gkw969. 10.1093/nar/gkw969.

20. Epshtein, V., Toulmé, F., Rahmouni, A.R., Borukhov, S., and Nudler, E. (2003). Transcription through the roadblocks: the role of RNA polymerase cooperation. EMBO J. 22, 4719–4727. 10.1093/emboj/cdg452.

21. Nudler, E. (2012). RNA Polymerase Backtracking in Gene Regulation and Genome Instability. Cell 149, 1438–1445. 10.1016/j.cell.2012.06.003.

22. Borukhov, S., Sagitov, V., and Goldfarb, A. (1993). Transcript cleavage factors from E. coli. Cell 72, 459–466. 10.1016/0092-8674(93)90121-6.

23. Fish, R.N., and Kane, C.M. (2002). Promoting elongation with transcript cleavage stimulatory factors. Biochim. Biophys. Acta BBA - Gene Struct. Expr. 1577, 287–307. 10.1016/S0167-4781(02)00459-1.

24. Fernández-Coll, L., Potrykus, K., Cashel, M., and Balsalobre, C. (2020). Mutational analysis of Escherichia coli GreA protein reveals new functional activity independent of antipause and lethal when overexpressed. Sci. Rep. 10, 16074. 10.1038/s41598-020-73069-1.

25. Marraffini, L.A., and Sontheimer, E.J. (2008). CRISPR Interference Limits Horizontal Gene Transfer in Staphylococci by Targeting DNA. Science 322, 1843–1845. 10.1126/science.1165771.

26. Laptenko, O., Lee, J., Lomakin, I., and Borukhov, S. (2003). Transcript cleavage factors GreA and GreB act as transient catalytic components of RNA polymerase. EMBO J. 22, 6322–6334. 10.1093/emboj/cdg610.

27. Deaconescu, A.M. (2021). Mfd – at the crossroads of bacterial DNA repair, transcriptional regulation and molecular evolvability. Transcription 12, 156–170. 10.1080/21541264.2021.1982628.

28. Gowrishankar, J., Leela, J.K., and Anupama, K. (2013). R-loops in bacterial transcription: Their causes and consequences. Transcription 4, 153–157. 10.4161/trns.25101.

29. Itaya, M., Omori, A., Kanaya, S., Crouch, R.J., Tanaka, T., and Kondo, K. (1999). Isolation of RNase H Genes That Are Essential for Growth of *Bacillus subtilis* 168. J. Bacteriol. 181, 2118– 2123. 10.1128/JB.181.7.2118-2123.1999.

30. Balibar, C.J., Shen, X., McGuire, D., Yu, D., McKenney, D., and Tao, J. (2010). cwrA, a gene that specifically responds to cell wall damage in Staphylococcus aureus. Microbiology 156, 1372–1383. 10.1099/mic.0.036129-0.

31. Reed, P., Atilano, M.L., Alves, R., Hoiczyk, E., Sher, X., Reichmann, N.T., Pereira, P.M., Roemer, T., Filipe, S.R., Pereira-Leal, J.B., et al. (2015). Staphylococcus aureus Survives with a Minimal Peptidoglycan Synthesis Machine but Sacrifices Virulence and Antibiotic Resistance. PLOS Pathog. 11, e1004891. 10.1371/journal.ppat.1004891.

32. Yan, J., Han, D., Liu, C., Gao, Y., Li, D., Liu, Y., and Yang, G. (2017). Staphylococcus aureus VraX specifically inhibits the classical pathway of complement by binding to C1q. Mol. Immunol. 88, 38–44. 10.1016/j.molimm.2017.05.018.

33. McAleese Fionnuala, Wu Shang Wei, Sieradzki Krzysztof, Dunman Paul, Murphy Ellen, Projan Steven, and Tomasz Alexander (2006). Overexpression of Genes of the Cell Wall Stimulon in Clinical Isolates of Staphylococcus aureus Exhibiting Vancomycin-Intermediate-S. aureus-Type Resistance to Vancomycin. J. Bacteriol. 188, 1120–1133. 10.1128/jb.188.3.1120-1133.2006.

34. Pyenson, N.C., Gayvert, K., Varble, A., Elemento, O., and Marraffini, L.A. (2017). Broad Targeting Specificity during Bacterial Type III CRISPR-Cas Immunity Constrains Viral Escape. Cell Host Microbe 22, 343–353.e3. 10.1016/j.chom.2017.07.016.

35. Yuzenkova, Y., Gamba, P., Herber, M., Attaiech, L., Shafeeq, S., Kuipers, O.P., Klumpp, S., Zenkin, N., and Veening, J.-W. (2014). Control of transcription elongation by GreA determines rate of gene expression in Streptococcus pneumoniae. Nucleic Acids Res. 42, 10987–10999. 10.1093/nar/gku790.

36. Goldberg, G.W., Jiang, W., Bikard, D., and Marraffini, L.A. (2014). Conditional tolerance of temperate phages via transcription-dependent CRISPR-Cas targeting. Nature 514, 633–637. 10.1038/nature13637.

37. Halpern, D., Chiapello, H., Schbath, S., Robin, S., Hennequet-Antier, C., Gruss, A., and El Karoui, M. (2007). Identification of DNA Motifs Implicated in Maintenance of Bacterial Core Genomes by Predictive Modeling. PLoS Genet. 3, e153. 10.1371/journal.pgen.0030153.

38. Yeeles, J.T.P., and Dillingham, M.S. (2010). The processing of double-stranded DNA breaks for recombinational repair by helicase–nuclease complexes. DNA Repair 9, 276–285. 10.1016/j.dnarep.2009.12.016.

39. Gopalkrishnan, S., Ross, W., Akbari, M.S., Li, X., Haycocks, J.R.J., Grainger, D.C., Court, D.L., and Gourse, R.L. (2022). Homologs of the Escherichia coli F Element Protein TraR, Including Phage Lambda Orf73, Directly Reprogram Host Transcription. mBio 13, e00952–22. 10.1128/mbio.00952-22.

40. Stepanova, E.V., Shevelev, A.B., Borukhov, S.I., and Severinov, K.V. (2009). Mechanisms of action of RNA polymerase-binding transcription factors that do not bind to DNA. Biophysics 54, 555–568. 10.1134/S0006350909050017.

41. Blankschien, M.D., Potrykus, K., Grace, E., Choudhary, A., Vinella, D., Cashel, M., and Herman, C. (2009). TraR, a Homolog of a RNAP Secondary Channel Interactor, Modulates Transcription. PLoS Genet. 5, e1000345. 10.1371/journal.pgen.1000345.

42. Camargo, A.P., Nayfach, S., Chen, I.-M.A., Palaniappan, K., Ratner, A., Chu, K., Ritter, S.J., Reddy, T.B.K., Mukherjee, S., Schulz, F., et al. (2023). IMG/VR v4: an expanded database of uncultivated virus genomes within a framework of extensive functional, taxonomic, and ecological metadata. Nucleic Acids Res. 51, D733–D743. 10.1093/nar/gkac1037.

43. Davidson, A.R., Lu, W.-T., Stanley, S.Y., Wang, J., Mejdani, M., Trost, C.N., Hicks, B.T., Lee, J., and Sontheimer, E.J. (2020). Anti-CRISPRs: Protein Inhibitors of CRISPR-Cas Systems. Annu. Rev. Biochem. 89, 309–332. 10.1146/annurev-biochem-011420-111224.

44. Sahakyan, H., Makarova, K.S., and Koonin, E.V. (2023). Search for Origins of Anti-CRISPR Proteins by Structure Comparison. CRISPR J. 6, 222–231. 10.1089/crispr.2023.0011.

45. Altschul, S.F., Gish, W., Miller, W., Myers, E.W., and Lipman, D.J. (1990). Basic local alignment search tool. J. Mol. Biol. 215, 403–410. 10.1016/S0022-2836(05)80360-2.

46. Chou-Zheng, L., and Hatoum-Aslan, A. (2022). Critical roles for ‘housekeeping’ nucleases in type III CRISPR-Cas immunity. eLife 11, e81897. 10.7554/eLife.81897.

47. Jia, N., and Patel, D.J. (2021). Structure-based functional mechanisms and biotechnology applications of anti-CRISPR proteins. Nat. Rev. Mol. Cell Biol. 22, 563–579. 10.1038/s41580-021-00371-9.

48. Chou-Zheng, L., Howell, O., Boyle, T.A., Hossain, M., Walker, F.C., Sheriff, E.K., Aslan, B., and Hatoum-Aslan, A. (2024). AcrIIIA1 is a protein–RNA anti-CRISPR complex that targets core Cas and accessory nucleases. Nucleic Acids Res. 52, 13490–13514. 10.1093/nar/gkae1006.

49. Zhu, M., Mu, H., Han, F., Wang, Q., and Dai, X. (2021). Quantitative analysis of asynchronous transcription-translation and transcription processivity in Bacillus subtilis under various growth conditions. iScience 24, 103333. 10.1016/j.isci.2021.103333.

50. Vogel, U., and Jensen, K.F. (1994). The RNA chain elongation rate in Escherichia coli depends on the growth rate. J. Bacteriol. 176, 2807–2813. 10.1128/jb.176.10.2807-2813.1994.

51. Sivaramakrishnan, P., Sepúlveda, L.A., Halliday, J.A., Liu, J., Núñez, M.A.B., Golding, I., Rosenberg, S.M., and Herman, C. (2017). The transcription fidelity factor GreA impedes DNA break repair. Nature 550, 214–218. 10.1038/nature23907.

52. Langmead, B., and Salzberg, S.L. (2012). Fast gapped-read alignment with Bowtie 2. Nat. Methods 9, 357–359. 10.1038/nmeth.1923.

53. Liao, Y., Smyth, G.K., and Shi, W. (2014). featureCounts: an efficient general purpose program for assigning sequence reads to genomic features. Bioinformatics 30, 923–930. 10.1093/bioinformatics/btt656.

54. Love, M.I., Huber, W., and Anders, S. (2014). Moderated estimation of fold change and dispersion for RNA-seq data with DESeq2. Genome Biol. 15, 550. 10.1186/s13059-014-0550-8.

55. Nayfach, S., Camargo, A.P., Schulz, F., Eloe-Fadrosh, E., Roux, S., and Kyrpides, N.C. (2021). CheckV assesses the quality and completeness of metagenome-assembled viral genomes. Nat. Biotechnol. 39, 578–585. 10.1038/s41587-020-00774-7.

56. Larralde, M. (2022). Pyrodigal: Python bindings and interface to Prodigal,an efficient method for gene prediction in prokaryotes. J. Open Source Softw. 7, 4296. 10.21105/joss.04296.

57. Hyatt, D., Chen, G.-L., LoCascio, P.F., Land, M.L., Larimer, F.W., and Hauser, L.J. (2010). Prodigal: prokaryotic gene recognition and translation initiation site identification. BMC Bioinformatics 11, 119. 10.1186/1471-2105-11-119.

58. Camargo, A.P., Roux, S., Schulz, F., Babinski, M., Xu, Y., Hu, B., Chain, P.S.G., Nayfach, S., and Kyrpides, N.C. (2024). Identification of mobile genetic elements with geNomad. Nat. Biotechnol. 42, 1303–1312. 10.1038/s41587-023-01953-y.

59. Kosmopoulos, J.C., Martin, C., Wainaina, J.M., Bolduc, B., Urvoy, M., Sullivan, M.B., and Anantharaman, K. (2026). CheckAMG: Accurate Identification of Auxiliary Viral Genes with Genome-Language Models. Preprint, 10.64898/2026.09.23.753886 https://doi.org/10.64898/2026.09.23.753886.

60. Larralde, M., and Zeller, G. (2023). PyHMMER: a Python library binding to HMMER for efficient sequence analysis. Bioinformatics 39, btad214. 10.1093/bioinformatics/btad214.

61. Eddy, S.R. (2011). Accelerated Profile HMM Searches. PLoS Comput. Biol. 7, e1002195. 10.1371/journal.pcbi.1002195.

62. Terzian, P., Olo Ndela, E., Galiez, C., Lossouarn, J., Pérez Bucio, R.E., Mom, R., Toussaint, A., Petit, M.-A., and Enault, F. (2021). PHROG: families of prokaryotic virus proteins clustered using remote homology. NAR Genomics Bioinforma. 3, lqab067. 10.1093/nargab/lqab067.

63. Kanehisa, M., Sato, Y., Kawashima, M., Furumichi, M., and Tanabe, M. (2016). KEGG as a reference resource for gene and protein annotation. Nucleic Acids Res. 44, D457–D462. 10.1093/nar/gkv1070.

64. Mistry, J., Chuguransky, S., Williams, L., Qureshi, M., Salazar, G.A., Sonnhammer, E.L.L., Tosatto, S.C.E., Paladin, L., Raj, S., Richardson, L.J., et al. (2021). Pfam: The protein families database in 2021. Nucleic Acids Res. 49, D412–D419. 10.1093/nar/gkaa913.

65. Zhou, K., Kosmopoulos, J.C., Colón, E.D., Badciong, P.J., and Anantharaman, K. (2026). V- and VL-scores unveil viral signatures and origins of protein families. Nat. Commun. 17, 5828. 10.1038/s41467-026-72028-0.

66. Abramson, J., Adler, J., Dunger, J., Evans, R., Green, T., Pritzel, A., Ronneberger, O., Willmore, L., Ballard, A.J., Bambrick, J., et al. (2024). Accurate structure prediction of biomolecular interactions with AlphaFold 3. Nature 630, 493–500. 10.1038/s41586-024-07487-w.

67. Katoh, K., and Standley, D.M. (2013). MAFFT Multiple Sequence Alignment Software Version 7: Improvements in Performance and Usability. Mol. Biol. Evol. 30, 772–780. 10.1093/molbev/mst010.

68. Wong, T.K.F., Ly-Trong, N., Ren, H., Demotte, P., Baños, H., Roger, A.J., Susko, E., Bielow, C., De Maio, N., Goldman, N., et al. (2026). IQ-TREE 3: phylogenomic inference software using complex evolutionary models. Mol. Biol. Evol. 43, msag117. 10.1093/molbev/msag117.

69. Kalyaanamoorthy, S., Minh, B.Q., Wong, T.K.F., Von Haeseler, A., and Jermiin, L.S. (2017). ModelFinder: fast model selection for accurate phylogenetic estimates. Nat. Methods 14, 587–589. 10.1038/nmeth.4285.

70. Russel, J., Pinilla-Redondo, R., Mayo-Muñoz, D., Shah, S.A., and Sørensen, S.J. (2020). CRISPRCasTyper: Automated Identification, Annotation, and Classification of CRISPR-Cas Loci. CRISPR J. 3, 462–469. 10.1089/crispr.2020.0059.

