## Supplemental Figures for "RNA Polymerase Backtracking Promotes Type III-A CRISPR DNA Targeting"

#### Figure S1

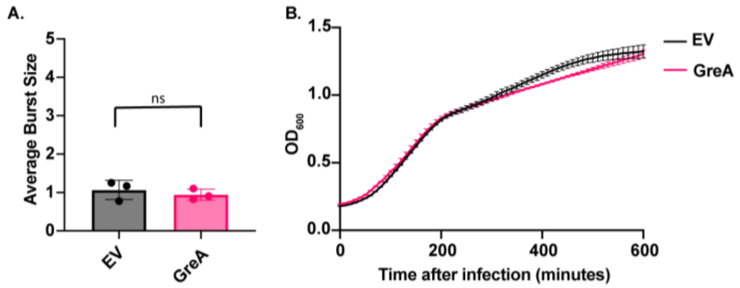


**Figure S1**. **GreA overexpression does not impact type II CRISPR (Cas9) anti-phage immunity**. **A.** Average burst size of ΦNM1γ6 in RN4220 carrying a type II-A CRISPR targeting the *gp12* gene of ΦNM1γ6 and a GreA overexpression plasmid, or an empty vector control (EV). Mean ± SD of 3 replicates is reported. **B.** Growth curve of RN4220 carrying the type II-A CRISPR targeting gp12 in the presence of ΦNM1γ6 at an MOI of 0.1. Cells also carry a GreA overexpression plasmid or the empty vector (EV). Mean ± SD of 3 replicates is reported. Statistics were performed with a unpair, two-sided t-test. ns: p>0.05; *: p<0.05; **: p<0.01; ***: p<0.001

#### Figure S2

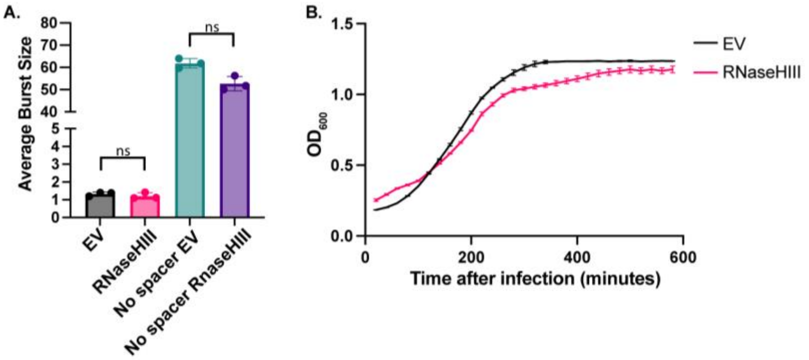


**Figure S2. Overexpression of RNaseHIII does not impact type III-A CRISPR anti-phage immunity.** **A.** Average burst size of ΦNM1γ6 in RN4220 carrying type III-A CRISPR with or without at targeting spacer against the *gp14* gene of ΦNM1γ6 and a RNaseHIII overexpression plasmid or the empty vector control (EV). Mean ± SD of 3 replicates is reported. **B.** Growth curve of RN4220 carrying the type II-A CRISPR targeting gp12 in the presence of ΦNM1γ6 at an MOI of 0.1. Cells also carry a RNaseHIII overexpression plasmid or the empty vector (EV). Mean ± SD of 3 replicates is reported. Statistics were performed with a unpair, two-sided t-test. ns: p>0.05; *: p<0.05; **: p<0.01; ***: p<0.001

**Figure S3**

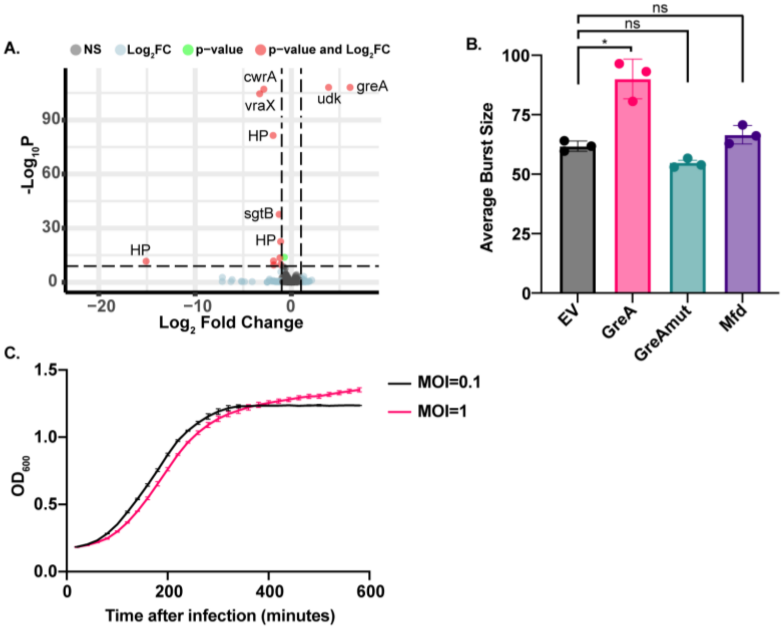


**Figure S3. Effects of GreA overexpression in the absence of type III-A CRISPR immunity.**
**A.** Differential gene expression analysis of RNA-sequencing results. The transcriptomes of RN4220 carrying the GreA overexpression plasmid were compared to that carrying the empty vector. Three biological replicates were evaluated for each condition. Horizontal dotted lines represent an adjusted p-value of 0.002, while vertical dotted lines represent a log2 fold-change of –2 or 2 in transcript levels. Transcripts with a log2 fold change between –2 or 2 and a p-value >0.002 are labeled as grey dots (Not significant, NS); transcripts that pass either the fold change or p-value cutoff (but not the other) are represented as blue and green dots, respectively; transcripts that pass both cutoffs are shown as red dots. Raw values are found in Supplemental Table 1. **B.** Average burst size of ΦNM1γ6 in RN4220 carrying type III-A CRISPR without a targeting spacer and either a GreA, GreAmut, or Mfd overexpression plasmid, as well as the empty vector (EV). Mean ± SD of 3 replicates is reported. **C.** Growth curve of RNA containing type III-A CRISPR targeting the phage gp14 gene in the presence of ΦNM1γ6 at an MOI of 0.1 or 1. Mean ± SD of 3 replicates is reported. Statistics were performed with an unpaired, two-sided t-test. ns: p>0.05; *: p<0.05; **: p<0.01; ***: p<0.001

### Figure S4

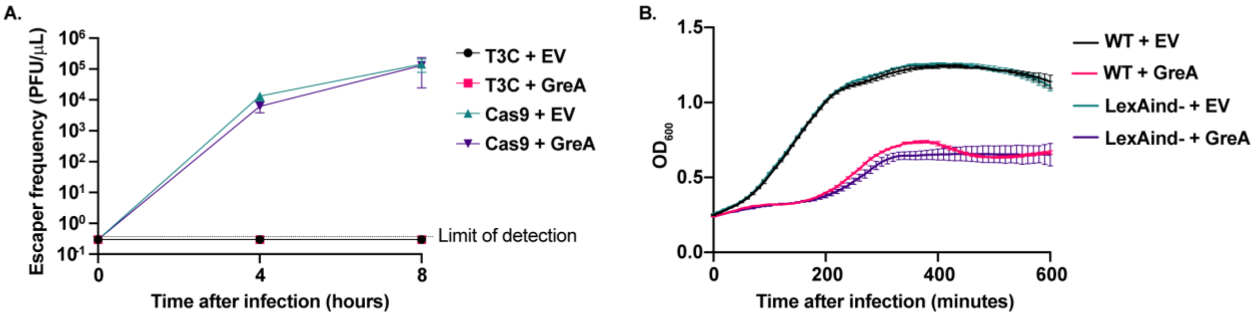


**Figure S4. Inhibition of type III-A CRISPR by GreA is independent of development of phage escaper and the SOS response. A.** Quantification of ΦNM1γ6 escaper mutants after infection of RN4220 cells carrying type III-A (T3C) or type II (Cas9) CRISPR systems targeting the *gp14* and *gp12* phage gene, respectively. Cells also carry a GreA overexpression plasmid or the empty vector (EV). Mean ± SD of 3 replicates is reported. **B.** Growth curve of RN4220 cells with an inducible (WT) or uninducible (LexAind-) SOS response. Strains are carrying the type III-A CRISPR targeting the *gp14* with the GreA overexpression plasmid or the empty vector (EV). Cells were infected with ΦNM1γ6 at an MOI of 0.1. Mean ± SD of 3 replicates is reported.

### Figure S5

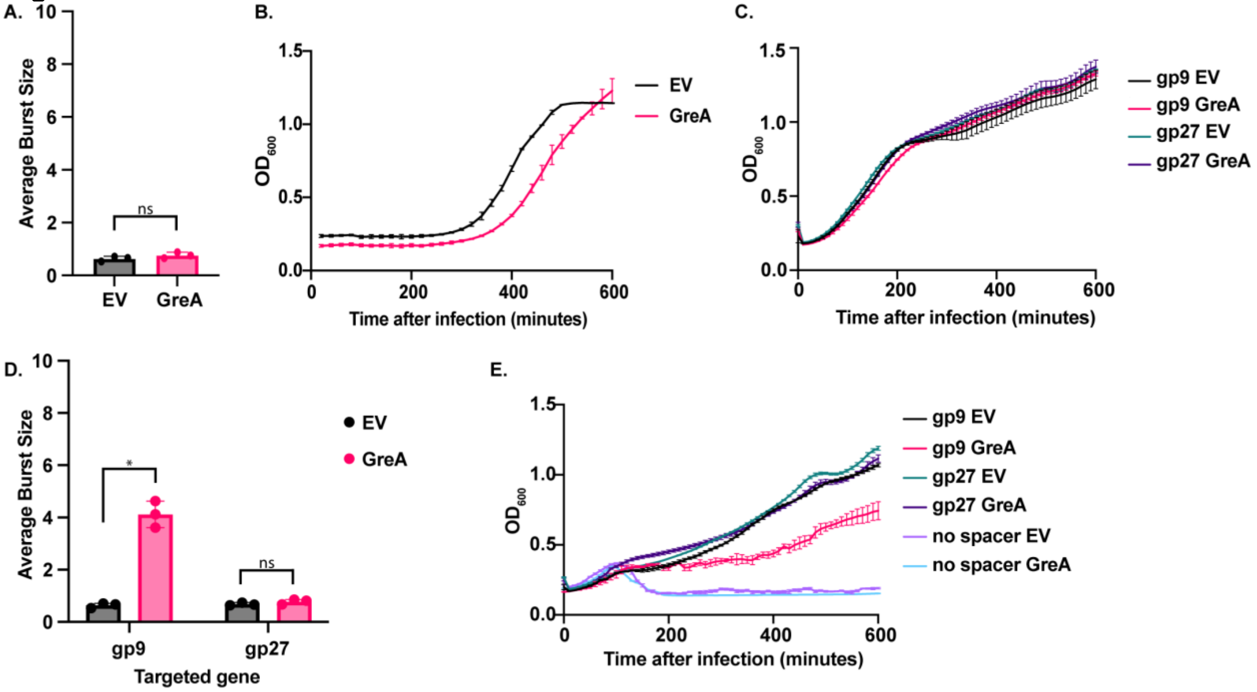


**Figure S5. Effect of GreA overexpression on type III-A CRISPR anti-phage defense when targeting phage late genes or phage Φ12. A.** Average burst size of ΦNM1γ6 in RN4220 with type III-A CRISPR targeting the phage *gp59* late gene and a GreA overexpression plasmid, or the empty vector (EV). Mean ± SD of 3 replicates is reported. **B.** Growth curve of RN4220 containing type III-A CRISPR targeting the phage gp59 gene in the presence of ΦNM1γ6 at an MOI of 0.1. Cells also contain a GreA overexpression plasmid, or the empty vector (EV). Mean ± SD of 3 replicates is reported. **C.** RN4220 growth curve containing type III-A CRISPR targeting the phage Φ12 gp9 early gene or gp27 late gene. Cells also contain a GreA overexpression plasmid, or the empty vector (EV). No phage was added. Mean ± SD of 3 replicates is reported. **D.** Average burst size of Φ12 in RN4220 containing type III-A CRISPR targeting the phage gp9 early gene or the gp27 late gene, and a GreA overexpression plasmid, or the empty vector (EV). Mean ± SD of 3 replicates is reported. **E.** RN4220 growth curve containing type III-A CRISPR targeting the phage gp9 early gene or the gp27 late gene in the presence of Φ12 at an MOI of 1. Cells also contain a GreA overexpression plasmid or the empty vector (EV). Mean ± SD of 3 replicates is reported. Where applicable, statistics were performed with an unpaired, two-sided t-test. ns: p>0.05; *: p<0.05; **: p<0.01; ***: p<0.001

**Figure S6**

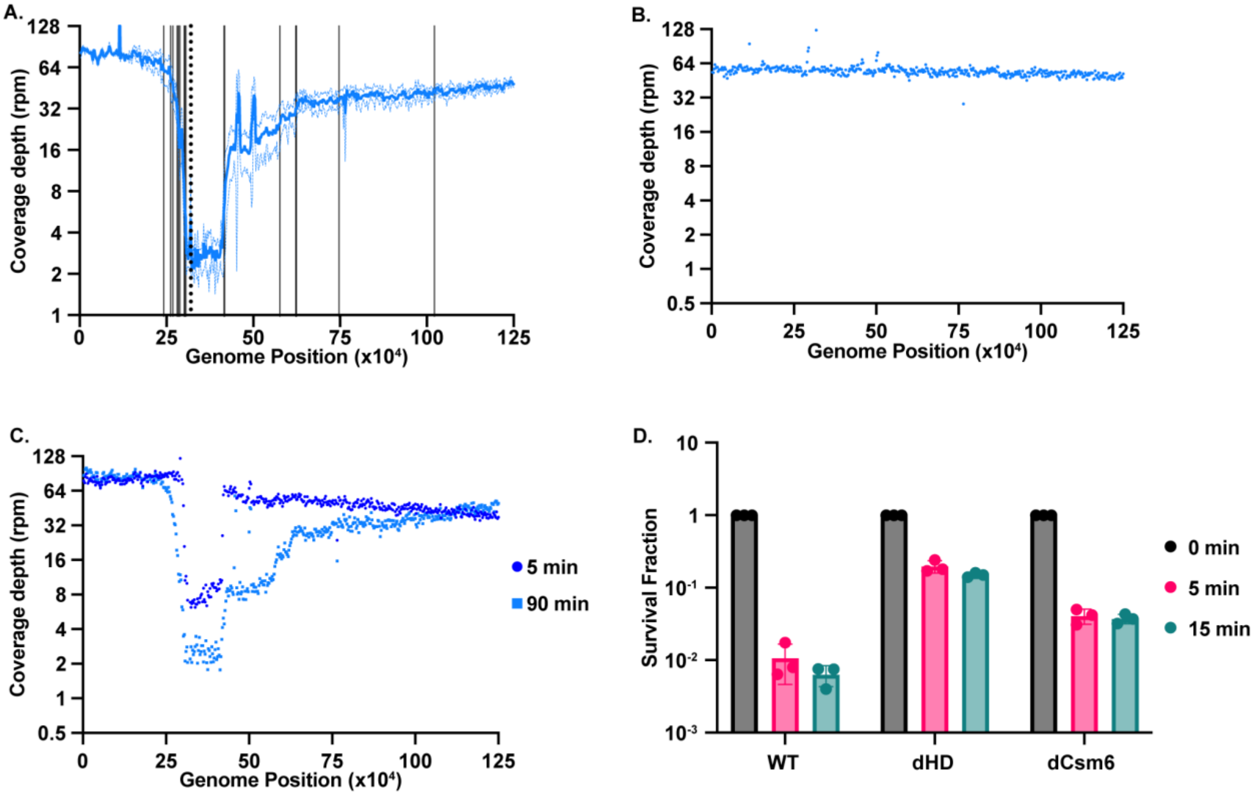


**Figure S6.** Type III-A CRISPR-induced DNA degradation in RN4220$_{Target}$ strain correlates with *chi* sites and is dependent on ssDNase activity of the HD domain. **A.** Coverage depth in reads per million (rpm) of RN4220$_{Target}$ containing type III-A CRISPR with a gp43 spacer 90 minutes after addition of aTc. Mean ± SD of 3 replicates is reported. Dotted vertical line shows the position of the inducible target, and solid vertical line shows position of *chi* sites in the genome. Shown are the first 125 kB of the RN4220 genome, with position 0 is set to the origin of replication. Marked are the chi sites upstream of the target from position 241,478 to 320,352 and downstream of the target from position 320,352 to 1,021,381. **B.** Coverage depth in reads per million (rpm) of RN4220$_{Target}$ containing a HD domain-deactivated type III-A CRISPR with a gp43 spacer 90 minutes after addition of aTc. **C.** Same as **B** but with cells containing a Csm6-deactivated type III-A CRISPR, and sequences were obtained 5 and 90 minutes after target induction. **D.** Survival fraction of RN4220$_{Target}$ containing a wild-type (WT), HD domain-deactivated (dHD), or Csm6-deactivated (dCsm6) type III-A CRISPR containing a gp43 spacer 5 and 15 minutes after aTc addition. Survival fraction is calculated by dividing the amount of CFU at the different time points by the amount of CFU before adding aTc (0 min). Mean ± SD of 3 replicates is reported.

**Figure S7**

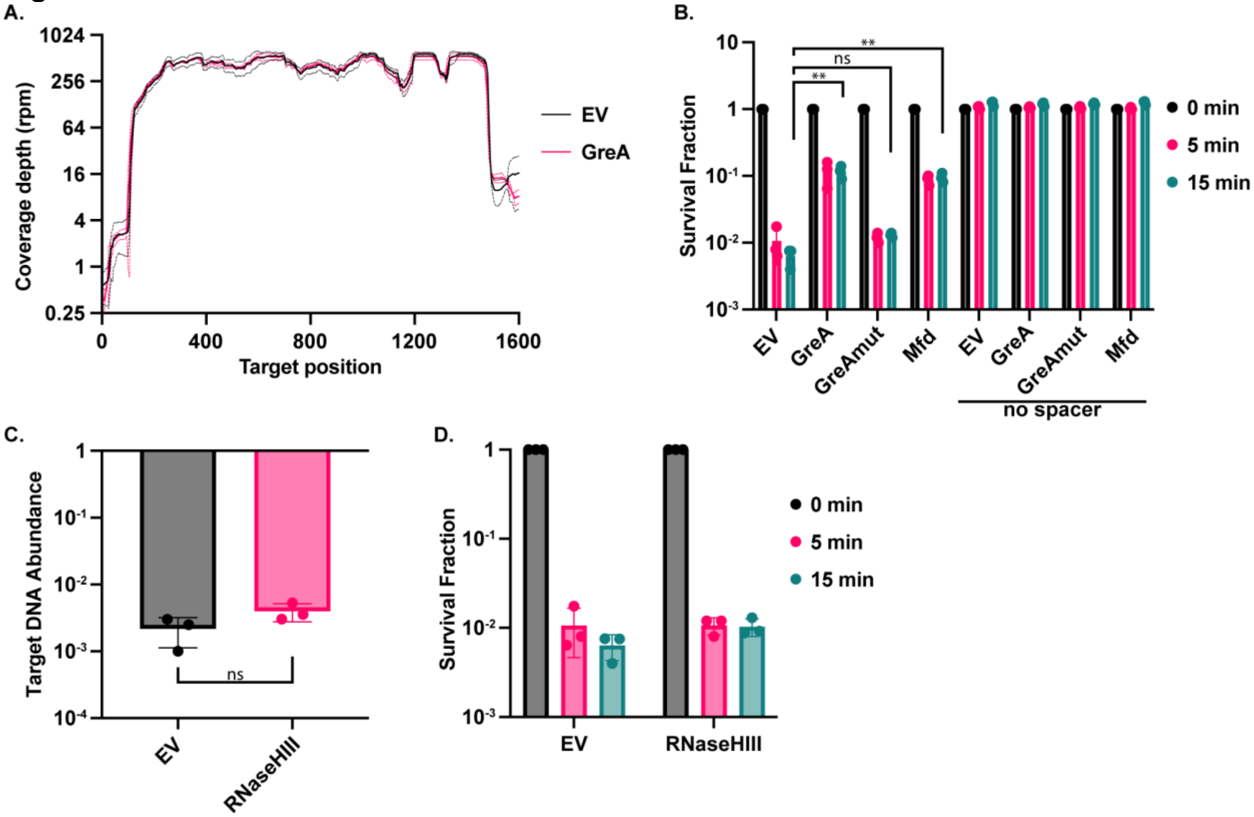


**Figure S7. Anti-backtracking factors inhibit type III-A CRISPR immunity in the RN4220$_{Target}$ strain without changing the target transcript levels. A.** RNA coverage depth in reads per million (rpm) of the inducible gp43 protospacer locus in RN4220$_{Target}$. Sequenced bacteria carries a type III-A CRISPR system without a targeting spacer and a GreA overexpression plasmid (or the empty vector, EV). Mean ± SD of 3 replicates is reported. **B.** Survival fraction of RN4220$_{Target}$ containing a type III-A CRISPR containing a gp43 spacer or without a spacer, and a GreA, GreAmut, or Mfd overexpression plasmid, or the empty vector (EV). Survival fraction is calculated by dividing the amount of CFU at the different time points after aTc addition by the amount of CFU before adding aTc (0 min). Mean ± SD of 3 replicates is reported. **C.** qPCR of the chromosomal gp43 protospacer in RN4220$_{Target}$ containing a type III-A CRISPR with a gp43 spacer 15 minutes after aTc addition. The cells also contained a RNaseHIII overexpression plasmid, or the empty vector (EV). Mean±SD of 3 replicates is shown. **D.** Same as **B** but with cells containing an RNaseHIII overexpression plasmid, or the empty vector (EV), and type III-A CRISPR with a gp43 spacer.

ns: $p>0.05$; *: $p<0.05$; **: $p<0.01$; ***: $p<0.001$

### Figure S8

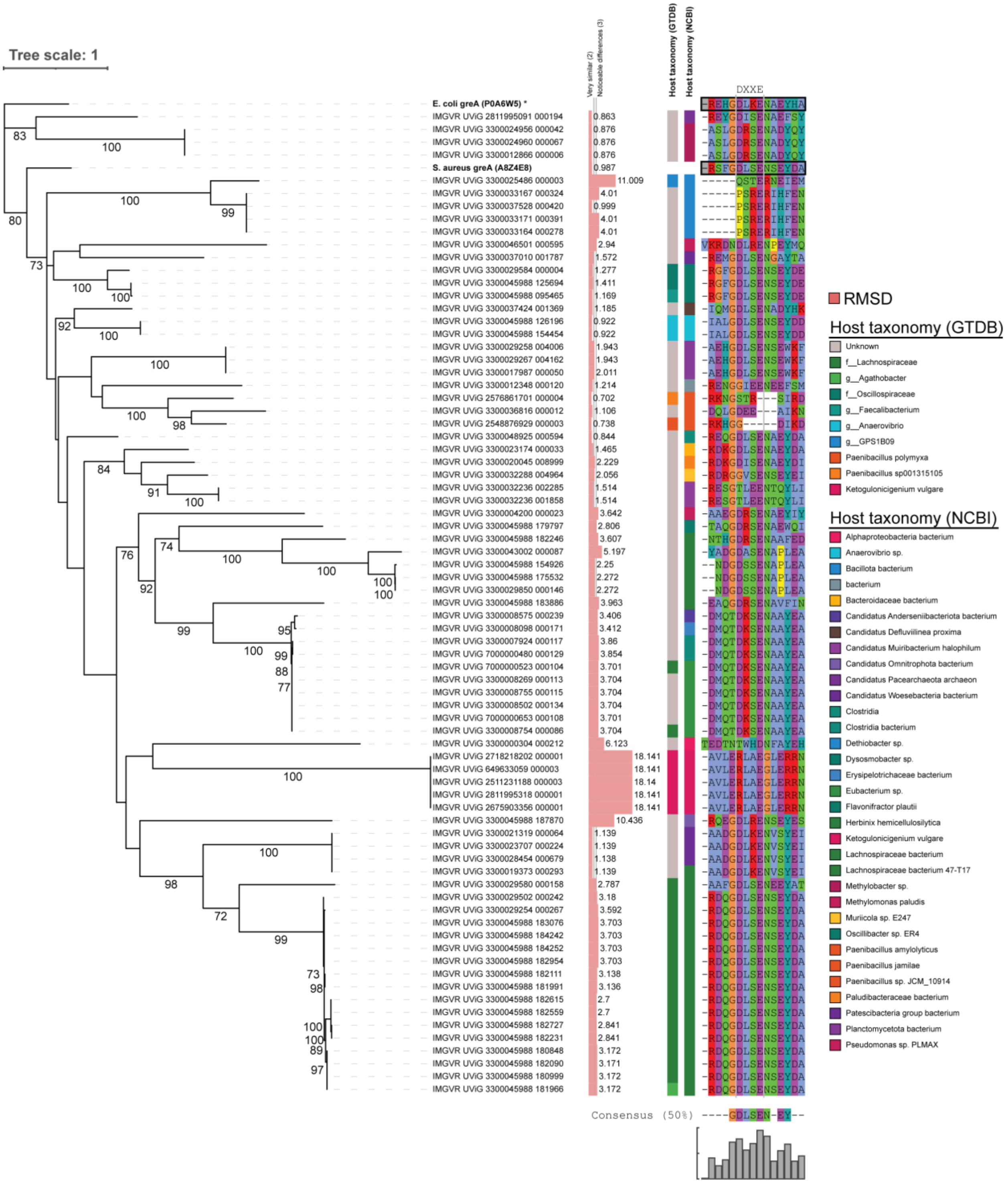


**Figure S8. Phylogeny of phage-encoded GreA-like sequences.** Phylogenetic tree of the 76 putative greA-like proteins found in the phage genomes, along with the *E. coli* (Uniprot: P0A6W5) and *S. aureus* (Uniprot: A8Z4E8) GreA bacterial proteins as reference. The gene sequences were aligned with MAFFT, and the tree was generated using IQ-Tree with ModelFinder, and 1000 bootstrap. Bootstrap values are shown in the node labels. Next to each branch are the RMSD values comparing the Alphafold3 predicted structure to the reference *E. coli* GreA, the host taxonomy according to IMG/VR (which uses GTDB) and according to the top match on NCBI nr, and finally the 87bp to 101 bp region in the aligned sequence to highlight the DXXE essential for anti-backtracking function motif in bacterial GreA. The Alphafold3 results, the sequence alignment file, the IQ-TREE3 output files, and the iTOL annotations are available at 10.5061/dryad.41ns1rnxk.

**Figure S9**

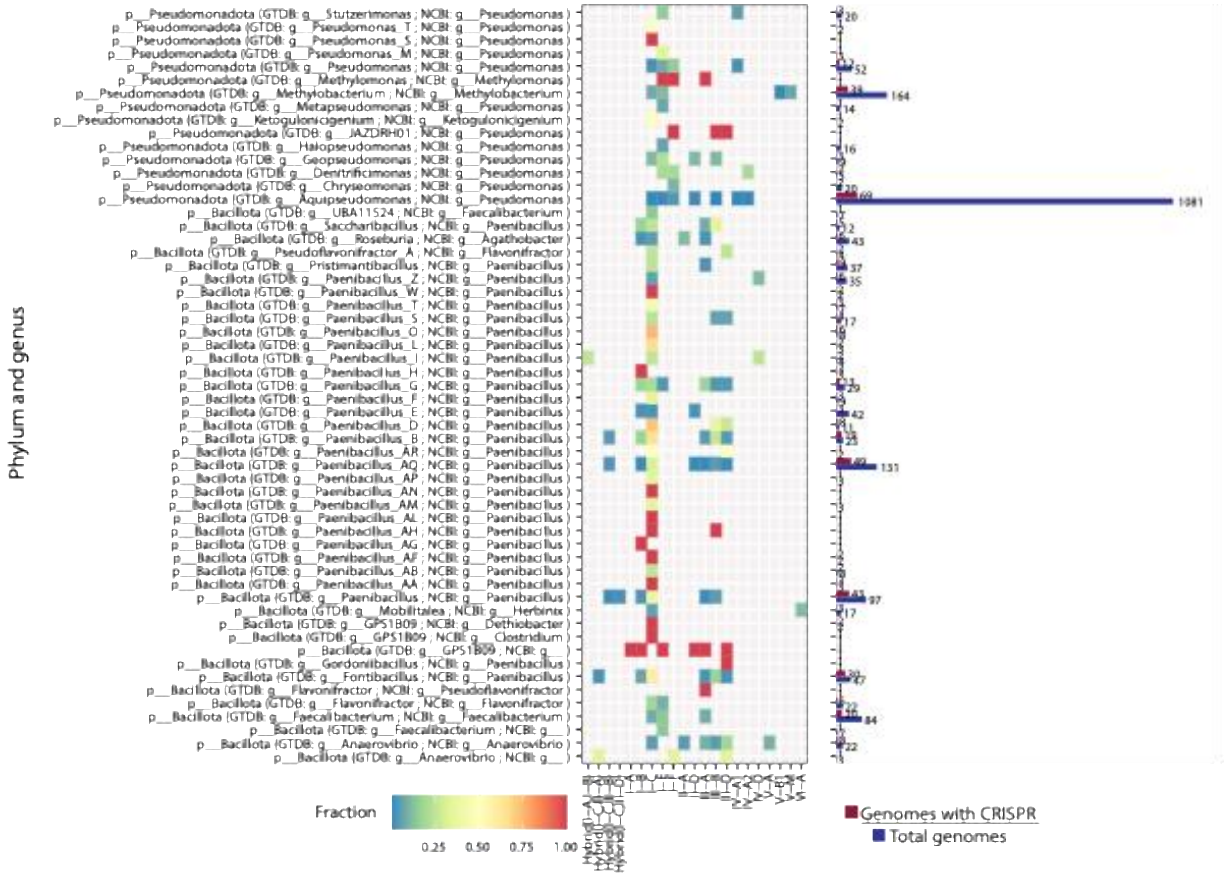


**Figure S9. Predicted hosts of the phages and CRISPR types present in the phage hosts.** We used cctyper to identify CRISPR systems in 16294 genomes retrieved using NCBI accession identifiers and obtained from the Genome Taxonomy Database (GTDB). The heatmap shows how many genomes have that type per total genomes in that taxonomic group. The bar plots show the total number of genomes with any identified CRISPR system and the total number of genomes in that genus that were originally analyzed.
